# An RNA-centric interactomics screen identifies novel proviral and antiviral genome binding interactors for tick-borne encephalitis virus

**DOI:** 10.64898/2026.04.23.720161

**Authors:** Axel Grot, Margaux Ligner, Cléa Torres, Yves Unterfinger, Véronique Legros, Muriel Coulpier, Guillaume Chevreux, Sandrine A. Lacour, Jennifer Richardson, Marion Sourisseau

## Abstract

The tick-borne encephalitis virus (TBEV), which belongs to the *Orthoflavivirus* genus, is a medically significant arbovirus in Europe and Asia. Despite distinct pathobiological outcomes, arboviruses have evolved specific strategies to co-opt cellular factors for replication and immune evasion, that involve both viral proteins and viral RNA. The RNA genome of TBEV, as both the substrate for genome replication and translation and a designated target for innate antiviral immunity, is a nexus for interactions with the proteome of host cells. Nonetheless, the molecular bases of these interactions and their implications for the replicative cycle of TBEV remain poorly understood. To create an inventory of such interactions and gain understanding of their functional import, we have resolved the set of cellular proteins bound to TBEV RNA in infected human cells using an agnostic RNA-centric approach.

Functional annotation of the resulting core interactome of 215 human host factors showed that viral RNA is deeply embedded in multiple cellular pathways, including those related to RNA and protein metabolism, cytoskeletal scaffolding, vesicle trafficking and innate antiviral immunity. For selected human interactors, we addressed their impact on TBEV infection in gene knockdown experiments, thereby identifying multiple restriction and dependency factors. These included sensors and effectors of innate immune pathways, as well as epitranscriptomic modifiers. Among the former, the dynamin-like GTPase MX2 protein, an interferon-stimulated gene with antiviral activity against multiple viruses including mosquito-borne orthoflaviviruses, displayed unexpected proviral activity against TBEV and a second tick-borne orthoflavivirus. Among the latter, WDR4, the non-catalytic component of the METTL1-WDR4 methyltransferase complex, emerged as a restriction factor with broad-spectrum activity against arboviruses belonging to multiple families of positive-strand RNA viruses.

In conclusion, this first description of the RNA interactome of a tick-borne orthoflavivirus illuminates the molecular interactions that underpin TBEV infection of human cells, which taken together reflect both the common ancestry of tick- and mosquito-borne orthoflaviviruses and their considerable evolutionary divergence.

## Introduction

Arboviruses form a polyphyletic group encompassing several viral families, among which *Flaviviridae* and *Togaviridae*, and include pathogenic viruses of major importance to human health, such as Japanese encephalitis virus (JEV, *Flaviviridae*) and Chikungunya virus (CHIKV, *Togaviridae*)^1,2^.

The most medically significant arbovirus in Europe is the tick-borne encephalitis virus (TBEV), a member of the *Orthoflavivirus* genus. TBEV is responsible for more than 3,000 annual infections that can induce severe encephalitis, causing death or significant neurological and neuropsychological sequelae. TBEV harbors an 11kb positive single-stranded RNA genome organized as a single open reading frame (ORF) flanked by two 5’ and 3’untranslated regions (UTR). Upon infection, this genome is translated into a polyprotein that is processed by viral and cellular proteases to yield 10 proteins, of which three are structural and seven non-structural (NS). The NS proteins assemble at the endoplasmic reticulum (ER) where they alter its structure, and together with poorly characterized recruited host factors, induce the formation of invaginated membrane structures called viral factories or replication organelles that constitute the site of viral replication. The newly synthesized viral RNA genome is encapsidated by the viral capsid (C) protein to form the ribonucleoprotein, and viral particles bud from the ER where they acquire their envelope. The newly formed virions then egress from infected cells through the exocytic pathway^3^.

As with any obligate intracellular parasite, the lifecycle of TBEV critically depends on the host cell machinery. Although armed with merely an 11kb RNA genome and ten proteins, the virus is capable of hijacking host resources by establishing interactions between its components and cellular proteins, leading to profound reprogramming of the host cell physiology and contributing to pathogenesis. The exact nature of the interactions underpinning this ability remains to be fully elucidated.

The development of high-throughput interactomic approaches has changed the way such interactions can be investigated and has recently contributed to the identification of novel host dependency and restriction factors for several orthoflaviviruses, providing insights into how they subvert host cell proteins to complete their lifecycles. These studies have helped uncover interactions that facilitate viral infection, either by promoting replication or by dampening the immune response, as well as host-driven interactions that act to restrict the infection. While most of them have focused on protein-protein interactions^4–10^ (PPI) between virus and host, some have addressed the viral RNA-host protein interactions (RPI), given the essential role of viral RNA in viral function and composition^11–16^. The viral genome indeed acts as a functional hub, recruiting host factors to promote its translation and replication^16–20^, while simultaneously being targeted by cellular antiviral immune sensors and effectors deployed to counter the infection^21–23^. While RPI for mosquito-borne orthoflaviviruses have been addressed in several studies, RPI of their tick-borne counterparts have only been addressed in a single publication. In this instance, mass spectrometry profiling, performed in a non-infectious context, was used to identify human proteins interacting with the TBEV genome 3’UTR, among which the fragile X messenger ribonucleoprotein (FMRP) emerged as a critical proviral factor^24^.

In this context, we have resolved the set of human proteins interacting with the TBEV genome within infected cells using comprehensive identification of RBPs by mass spectrometry (ChIRP-MS) methodology^25^. By gene silencing for a subset of these interactors, we have uncovered the role of multiple host factors, whether post-transcriptional modifiers, innate immune effectors and sensors, or mRNA decay effectors, that display significant proviral or antiviral activity during the replication cycle of TBEV. We subsequently assessed their impact on other arboviruses, notably revealing a broadly conserved antiviral phenotype for the WDR4 protein, a component of the heteromeric METTL1-WDR4 epitranscriptomic modifier.

## Materials and methods

### Cell culture

The human epithelial cell line A549, used to resolve the TBEV RNA interactome as well as to perform RNA interference assays, the simian epithelial cell line Vero, used for propagation and titration of TBEV, and the human kidney epithelial-like cell line 293T were all cultivated in DMEM (Dulbecco/Vogt modified Eagle’s minimal essential medium, Gibco, Waltham, USA) supplemented with 5% fetal bovine serum (FBS) (Eurobio, Les Ulis, France).

### Viral propagation and quantification

The TBEV Hypr strain (GenBank accession number U39292.1), originally isolated in the Czech Republic in 1953 from a child with tick-borne encephalitis, was kindly provided by Sara Moutailler (UMR BIPAR, Anses, France). The Louping Ill virus (LIV) LI3/1 strain (GenBank accession number KP144331.1) was kindly provided by Nicholas Johnson (Animal and Plant Health Agency, United Kingdom). Both the CHIKV strain, which originated from an outbreak in Réunion Island in 2005 (GenBank accession number AM258990.1), and the JEV Nakayama strain (GenBank accession number EF571853.1) were kindly provided by the European reference Laboratory for Equine Diseases (Anses, France).

Viral stocks were prepared as described previously^9^, and quantified by endpoint dilution assay on Vero cells, as visualized by the virus-induced cytopathic effect (CPE). Briefly, 1.5 × 10^4^ Vero cells were seeded in 100 μL of DMEM and 5% FBS in wells of a 96-well plate the day prior to infection with 30 μL of serial dilutions of virus in medium as above (6 wells per dilution). Five according to the method of Reed and Muench^26^.

### TBEV RT-qPCR

RNA was extracted from cell lysate derived from ChIRP-MS samples using TRizol (Invitrogen, Waltham, USA) and eluted in 30 µl of RNAse-free H_2_O (Qiagen, Venlo, Netherlands) For quantifying TBEV RNA, quantitative reverse transcriptase PCR (RT-qPCR) was performed on 2 µl of extracted RNA using primers specific for the envelope gene (E) of TBEV Hypr (5′ TGT TTC CAT GGC AGA GCC AG and 5′ TCC TTG AGC TTG ACA AGA CAG) and probe (5′ 6-carboxyfluorescein [FAM]- T GGA ACA CCT TCC AAC GGC TTG GCA-minor groove binder nonfluorescent quencher [MGB-NFQ]) (Eurobio, Les Ulis, France) and the qScript XLT 1-Step RT-qPCR ToughMix (Quantabio, Beverly, USA) using the LightCycler 96 real-time PCR system (Roche Life Science, Basel, Switzerland).

Quantification of TBEV RNA copies was performed in reference to a standard curve of TBEV Hypr E RNA that had been *in vitro*-transcribed from a pGEM plasmid (Promega Corporation, Madison, USA) containing the TBEV Hypr E ORF, using the RiboMAX™ Express Large Scale T7 RNA production system (Promega).

### Comprehensive Identification of RBPs by Mass Spectrometry (ChIRP-MS)

The oligonucleotide probes used for oligoprecipitation were designed using the Stellaris Probe Designer tool (https://www.biosearchtech.com/support/tools/design-software/stellaris-probe-designer). The TBEV Hypr genome sequence was submitted with parameters set to generate 50 probes of 20 nucleotides in length, covering the entire viral genome at intervals of approximately 200 nucleotides, using masking level 2. The resulting probe sequences are listed in Supplementary Table 2. The probes were modified with a 3’-biotin-TEG (triethylene glycol) motif.

To perform ChIRP-MS on the human A549 cell line, each condition—that is, uninfected (NI) and infected (TBEV)—consisted of three replicates, each comprising approximately 10^8^ cells on the day of infection. At day 0, cells in the infected condition were inoculated with TBEV Hypr at an MOI of 5.60 × 10⁻⁵. Uninfected (NI) control cells received the same volume of a mock inoculum.

Forty-eight hours post-infection, cells were detached and centrifuged at 515 × g for 5 minutes. Supernatants were discarded, and cell pellets were weighed. They were then crosslinked using 4% formaldehyde under agitation at room temperature for 30 minutes. The formaldehyde was neutralized by adding 2M glycine to each tube, followed by 5 minutes of agitation at room temperature. Tubes were centrifuged as above.

The resulting pellets were then solubilized In 1 mL of lysis buffer (50 mM Tris-HCl pH 7.0, 10 mM EDTA, 1% SDS, supplemented with rNase inhibitors (SuperaseIn, Gibco) per 100 mg of cells. Lysates were sonicated using a Hielscher UP200St sonicator (Teltow, Germany), set to cycles of 30 seconds ON / 45 seconds OFF at 20W power, for 35 minutes.

After sonication, 2mL of hybridization buffer (750 mM NaCl, 1% SDS, 50 mM Tris-HCl pH 7.0, 1 mM EDTA, 15% formamide, supplemented with RNase and protease inhibitors (cOmplete, Roche) per 1mL of lysate were added to each tube. To remove nonspecific interactions, 50 µL of streptavidin-coated magnetic beads (Dynabeads MyOne C1, Invitrogen) were added per milliliter of lysate to each replicate. Tubes were gently vortexed, then rotated for 30 minutes at 37°C. Beads were separated from the supernatants using a magnetic rack; supernatants were transferred to new tubes, and the process was repeated to ensure complete removal of magnetic beads. To capture the complexes, 1.5 µL of a pool of 100 µM biotinylated oligonucleotides complementary to the TBEV genome were added per 1mL of lysate. Samples were gently vortexed and rotated overnight at 37°C to allow hybridization of the biotinylated probes to their targets.

After capture, 150 µL of pre-washed magnetic beads per 1mL of lysate were added to the tubes, which were then rotated at 37°C for 45 minutes. The captured complexes were precipitated using a magnetic rack and the beads were washed three times in 500 µL of wash buffer (2x SSC buffer, 0.5% SDS), each time rotating the tubes at 37°C for 10 minutes. Beads were then washed twice with 500 µL of a second wash buffer (50 mM Tris-HCl, 150 mM NaCl) following the same procedure. Finally, beads were washed once with RNase-free water and resuspended in RNase-free water. The beads were sent to the proteomics platform ProteoSeine (Institut Jacques Monod Paris, France) for mass spectrometry analysis.

To assess the efficiency of oligoprecipitation, a portion of magnetic beads was solubilized in 200 µL elution buffer (12.5 mM biotin, 7.5 mM HEPES, pH 7.9, 75 mM NaCl, 1.5 mM EDTA, 0.15% SDS, 0.075% sarkosyl, and 0.02% Na-deoxycholate) and rotated at room temperature for 20 minutes, then at 65°C for 15 minutes with manual shaking every minute. After separation, the eluates were collected on a magnetic rack. This step was repeated twice. To concentrate the proteins, 25µL of trichloroacetic acid was added per 100µL of eluate, vortexed vigorously and incubated overnight at 4°C. The next day, the eluates were centrifuged at 21,000 × g for 45 minutes at 4°C to precipitate the proteins. The supernatants were discarded and the precipitates washed in 1 mL of glacial acetone followed by centrifugation at 21,000 × g for 5 minutes at 4°C. The supernatants were discarded. A second centrifugation was performed to remove residual acetone. Precipitates were solubilized in 30 µL of 1x lithium dodecyl sulfate (LDS) (Invitrogen) supplemented with a reducing agent (Sample Reducing Agent 10X, Invitrogen). 15 µL of ChIRP-MS lysate were collected prior to oligoprecipitation and mixed with 15 µL of 2x LDS supplemented with reducing agent. All samples were then heated at 95°C for 10 minutes to denature proteins. The migration of all samples was performed on a 4-12% Bis-Tris polyacrylamide gel (Invitrogen). After migration, the gel was silver stained using the SilverQuest Staining Kit (Invitrogen) according to the manufacturer’s protocol. After staining, the gels were digitally photographed.

### LC-MS/MS analysis

MS-grade acetonitrile (ACN), H_2_O and formic acid (FA) were from ThermoFisher Scientific (Waltham, MA, USA). Sequencing-grade trypsin/Lys C mix was from Promega. Ammonium bicarbonate (NH_4_HCO_3_) was from Sigma-Aldrich (Saint-Louis, MO, USA).

Prior to LC-MS/MS analysis, beads from pulldown experiments were incubated overnight at 37°C with 20 μL of 50 mM NH_4_HCO_3_ buffer containing 1 µg of sequencing-grade trypsin/Lys C mix. The digested peptides were then desalted on evotips provided by Evosep (Odense, Denmark) according to the manufacturer’s procedure.

Samples were analyzed on a timsTOF Pro 2 mass spectrometer (Bruker Daltonics, Bremen, Germany) coupled to an Evosep one system (Evosep) operating with the 30SPD method developed by the manufacturer. Briefly, the method is based on a 44-min gradient and a total cycle time of 48 min with a C18 analytical column (0.15 x 150 mm, 1.9 µm beads, ref EV-1106) equilibrated at 40°C and operated at a flow rate of 500 nL/min. H_2_O/0.1 % FA was used as solvent A and ACN/ 0.1 % FA as solvent B. The timsTOF Pro 2 was operated in DDA-PASEF mode over a 1.3 sec cycle time^27^. Mass spectra for MS and MS/MS scans were recorded between 100 and 1700 m/z.

MS raw files were processed using PEAKS Online 12 (build 2.1, Bioinformatics Solutions Inc.). Data were searched against the *Homo sapiens* Swiss-Prot database with 11 additional sequences from the TBEV viruses (downloaded 2025_01, 20,656 entries). Parent mass tolerance was set to 20 ppm, with fragment mass tolerance at 0.05 Da. Semi-specific tryptic cleavage was selected and a maximum of 2 missed cleavages was authorized. For identification, the following post-translational modifications were included: oxidation (M), deamidation (NQ), acetylation (Protein N-term) and half of a disulfide bridge (C) as variables. Identifications were filtered based on a 1% FDR (false discovery rate) threshold at both peptide and protein group levels. Label-free quantification was performed using the PEAKS Online quantification module, allowing a mass tolerance of 1 ppm, a CCS error tolerance of 0.02 and a -0.25 min retention time shift tolerance for match between runs. Protein abundance was inferred using the Top-N method and TIC was used for normalization. Multivariate statistics on proteins and peptides were performed using Qlucore Omics Explorer 3.8 (Qlucore AB, Lund, SWEDEN). A positive threshold value of 1 was specified to enable a log2 transformation of abundance data for normalization; i.e., all abundance data values below the threshold are replaced by 1 before transformation. The transformed data were finally used for statistical analysis; i.e., evaluation of differentially represented proteins or peptides between two groups using a Student’s bilateral t-test. An adjusted p-value less than 0.05, as determined according to the Benjamini–Hochberg procedure (q-value), was used to filter differential candidates.

To distinguish specific interactors of the TBEV genome from nonspecific background, the list was filtered based on mass spectrometry analysis parameters. A minimum of two unique peptides was required for each protein. The peptide abundance ratio between the NI and TBEV conditions had to be at least two. The p-values associated with peptide abundance for each protein across TBEV replicates were set at a maximum of 5%, defining the extended interactome of the TBEV genome. By setting the p-values to a maximum of 1%, the core interactome of the TBEV genome was defined.

### GO Term enrichment analysis

Functional enrichment analysis of the extended and core interactome was achieved using R (v. 4.5.2) through RStudio (v. 2026.1.1.403) with the clusterProfiler package (v4.18.4) and with semantic similarity among GO terms computed with the GOSEmSim package (v2.36.0)^28,29^. Dendrograms were generated using the enrichplot package (v1.30.5)^29^. Comparison of functional enrichment between our dataset and previous ChIRP-MS studies was performed using Metascape^30^.

### Manual annotation

The 215 proteins from the core interactome were manually annotated based on scientific literature. Various features were used to describe each interactor: a previously reported impact on a viral life cycle (regardless of the virus) and if so the nature of impact; whether known to bind RNA and if so its functional role in that capacity; associated biological functions, cellular pathways, and cellular compartments; and whether it previously retrieved in other ChIRP-MS studies of orthoflaviviruses.

### Functional screening by RNAi

A549 cells were seeded in 48-well plates at a density of 1 × 10⁴ cells per well 72 hours prior to siRNA transfection. Cells were transfected in triplicate with a pool of four siRNAs per target of interest, or with a control siRNA at a final concentration of 20 µM using DharmaFECT 1 transfection reagent. The list of siRNAs is provided in Table S6.

Twenty-four hours post-transfection, the transfection medium was removed and replaced with fresh medium. Forty-eight hours post-transfection, cells were inoculated with viral stocks at a MOI of 1.11 × 10⁻⁴ for TBEV, 5.57 × 10⁻⁵ for LIV, 4.7 × 10⁻³ for CHIKV, and 1.58 × 10⁻³ for JEV. After 2 hours of inoculation, the viral inocula were removed and cells were washed three times with PBS. Forty-eight hours post-infection, the culture supernatants from all wells were collected and titrated by endpoint dilution.

### Cell viability assay

A total of 10,000 A549 cells per well were seeded in a 48-well plate, and cells in triplicate wells were transfected 72h later with siRNAs targeting the tested genes or with the NT siRNAs, following the same procedure as previously described; one triplicate of cells was left untransfected. After 48 hours of silencing, cell viability was assessed using the CellTiter-Glo® assay kit (Promega) as per the manufacturer’s recommendation. This method is based on ATP quantification, and the luminescence signal measured is proportional to cell viability. To determine the percentage of viability, all luminescence values were normalized to those measured in cells transfected with control siRNA NT.

### WDR4 KO A549 cell line generation

A A549-derived cell line knocked-out for WDR4 expression (WDR4-KO) was generated following the TrueGuide™ Synthetic gRNA procedure from Invitrogen. Briefly, 80 000 A549 cells were seeded in a 24-well plate and transfected 24h later with a mix containing the recombinant TrueCut™ Cas9 Protein v2 (Invitrogen), the TrueGuide™ Synthetic gRNA (Invitrogen) directed against the WDR4 gene, and the Lipofectamine™ CRISPRMAX™ (Invitrogen) transfection reagent. Cells were incubated at 37°C for 48h, then amplified and plated at 1, 0.1 and 0.01 cells per well in 96-well plates. Cell growth was visually monitored for several days to detect wells where a single colony could be observed. For such wells, cells were amplified, and the expression of the WDR4 protein was assessed by Western blot.

### Infection of KO-WDR4 A549-derived cells

To determine the impact of the WDR4 KO on the cycle of TBEV, wildtype (WT), WDR4-KO, WDR4-KO GFP transfected, and WDR4-KO rescued A549 cells were seeded in 24-well plates at a density of 100 000 cells per well 24 hours prior to TBEV infection. Cells were then inoculated with TBEV at MOI of 0.01 or 0.1. After 2h hours of inoculation, the viral inocula were removed and cells were washed three times with PBS. Forty-eight hours post-infection, the culture supernatants from all wells were collected and titrated by endpoint dilution. Infectious titers for each sample were calculated using the Reed & Muench method and expressed in TCID₅₀/mL of viral suspension.

### Construction of lentiviral vectors

WDR4 and GFP were expressed via lentiviral transduction using pLX301, a 3rd generation lentiviral provirus that utilizes an internal CMV promoter to drive transgene expression. The coding sequences of WDR4 and GFP, flanked by Gateway extension sites and a Flag tag in 3’termini, were amplified by PCR using oligonucleotides listed in Table S1, and WDR4_pCSdest and pT7-EGFP-C1-HsNot1 plasmids as templates, respectively. Each construct was designed to include a C-terminal FLAG epitope tag. The amplified products were first cloned into the pDONR207 entry vector (Invitrogen) and subsequently transferred into the pLX301, a 3rd generation lentiviral Gateway destination vector allowing puromycin selection, using Gateway® cloning technology (Invitrogen), following the manufacturer’s protocol. All PCR-amplified sequences and cloning junctions were verified by sequence analysis (Eurofin Genomics, Luxembourg, Luxembourg)

WDR4_pCSdest was a gift from Roger Reeves (Addgene plasmid #53953; http://n2t.net/addgene:53953 ; RRID:Addgene_53953)^31^. pT7-EGFP-C1-HsNot1 was a gift from Elisa Izaurralde (Addgene plasmid #37370 ; http://n2t.net/addgene:37370; RRID:Addgene_37370)^32^. pLX301 was a gift from David Root (Addgene plasmid #25895; http://n2t.net/addgene:25895 ; RRID:Addgene_25895)^33^.

### Lentiviral pseudoparticle generation and transduction

Pseudoparticles were generated by cotransfecting cells with three plasmids encoding (i) a provirus carrying the transgene or reporter of interest (pLX301-WDR4 or GFP), (ii) psPAX2, a 2nd generation lentiviral packaging plasmid, and (iii) pMD2.G, a VSV-G envelope expressing plasmid. 293T cells were seeded at 1.8 × 10⁶ cells per well in a six-well plate coated with poly-L-lysine (Sigma). Transfection was performed the next day with TransIT-LT1 transfection reagent (Mirus, Madison, WI) as per the manufacturer’s instructions. Supernatants were collected at 48 hours post-transfection and filtered through a 0.45-μm pore-size filter.

For lentiviral transduction, WDR4-KO A549 cells were seeded in a 12-well plate. The next day, cells were inoculated with 25μL of pseudoparticle preparation in complete medium for 24h. To allow recovery from transduction and stabilization of transgene expression, cells were passaged into progressively larger culture vessels over 1–2 weeks under puromycin selection (final concentration: 2µg/mL). Transgene expression was confirmed by immunoblotting.

psPAX2 (Addgene plasmid #12260; http://n2t.net/addgene:12260 ; RRID:Addgene_12260) and pMD2.G (Addgene plasmid #12259 ; http://n2t.net/addgene:12259; RRID:Addgene_12259) were gifts from Didier Trono.

### Immunoblot

For immunoblot analysis of WDR4 or GFP expression, 1 × 10^7^ cells were lysed in 1mL of RIPA buffer for 10 minutes on ice, followed by centrifugation to remove insoluble material. Protein concentration was determined using a BCA protein assay kit (Pierce). 5 μg of total protein from each sample was mixed with a one-quarter volume of 4× LDS sample buffer (Invitrogen) and reducing agent (Invitrogen), then heated at 95°C for 10 minutes.

Proteins were separated and analyzed by immunoblotting using anti-WDR4 (rabbit monoclonal – EPR11052, Abcam, Cambridge, UK), anti-GFP (mouse monoclonal – clones 7.1 and 13.1, Roche Life Science, Basel, Switzerland), and anti-GAPDH (rabbit monoclonal – clone 14C10, Ozyme, Saint-Cyr-l’École, France) antibodies. Horseradish peroxidase (HRP)-conjugated goat anti-mouse or goat anti-rabbit secondary antibodies (polyclonal – respectively 5178-2504 and 5196-2504, Bio-Rad, Hercules, USA) were used, and signal detection was performed using Clarity ECL Western Blotting (Bio-Rad) and the ChemiDoc imaging system (Bio-Rad).

### Statistical analysis

The p-values presented in the graphs of Figures 4, 5 and 6 and Supplementary Figure S3 were obtained using the non-parametric Kruskal–Wallis test, followed by Dunn’s multiple-comparison *post hoc* test with Benjamini–Krieger–Yekutieli two-stage step-up correction for multiple testing. For TBEV, each target siRNA was compared with the corresponding control siRNA NT across the three biological replicates per condition. For LIV, JEV and CHIKV, each target siRNA was compared with the corresponding target for TBEV. These analyses were performed using GraphPad Prism 10 (GraphPad Software, Boston, USA).

## Results

### The TBEV – Homo sapiens RNA-protein interaction network

To resolve the viral-RNA interactome within the natural context of TBEV infection in human cells, we used an unbiased RNA-centric approach known as ChIRP-MS (Figure 1A). Briefly, A549 cells were infected with the highly virulent TBEV Hypr strain or mock infected. Forty-eight hours after infection, when each individual replicate contained 108 to 109 copies of the RNA genome per 50 million cells (Figure 1B), infected (TBEV) and non-infected (NI) cultures were treated with formaldehyde to cross-link RNA to adjacent RNA-binding protein (RBP) complexes. Viral RNA-protein complexes were then specifically hybridized to 50 biotinylated oligonucleotide probes complementary to the full-length viral genome (Table S1), followed by streptavidin-coupled bead pulldown. Co-precipitated proteins were then identified by LC-MS/MS and bioinformatic analysis. Specific enrichment was controlled by elution of a subset of ribonucleoprotein (RNP) complexes captured on beads, which were run on SDS-PAGE and visualized by silver staining (Figure 1C).

**Figure 1.**
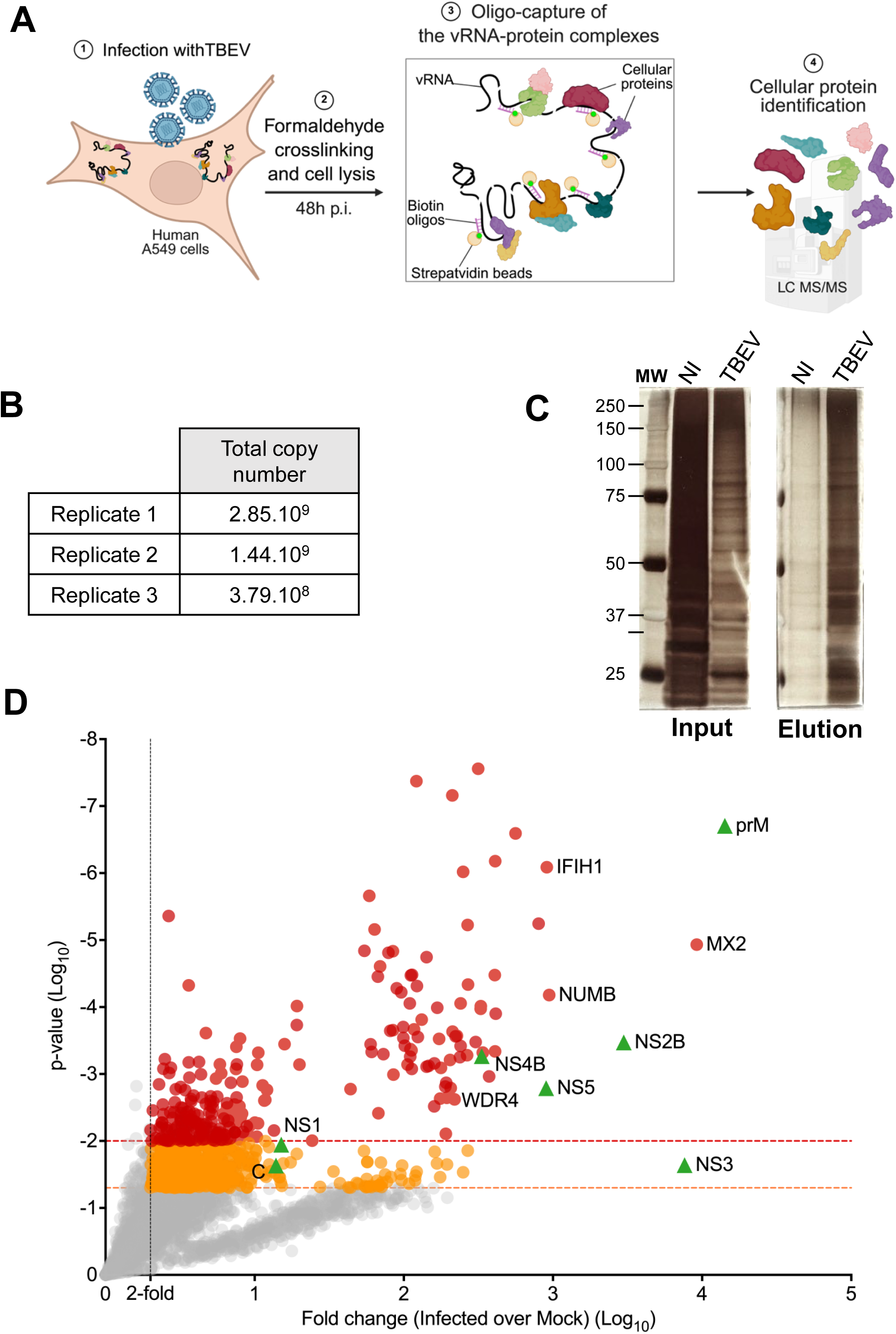
ChIRP-MS uncovers the TBEV genome protein interactome. (A) Schematic of the ChIRP-MS experimental procedure. (B) Quantification of TBEV RNA genome copies in each biological replicate by RT-qPCR prior to ChIRP-MS processing. (C) Visualization of protein enrichment in TBEV-infected versus mock-infected cells before and after the ChIRP procedure. 0.1% of each sample was resolved by SDS-PAGE and visualized by silver staining. (D) Scatter plot showing the enrichment ratio of TBEV-infected over uninfected samples for all proteins identified by ChIRP-MS, plotted against statistical significance. The ratios were calculated from the mean abundance of total peptides detected by mass spectrometry. Proteins specifically enriched on the TBEV genome exhibit an enrichment ratio greater than 2 with a p-value below 0.05. Proteins specifically interacting with the TBEV genome display an enrichment ratio greater than 2 with a p-value below 0.05. Orange and red dots highlight proteins belonging to the extended interactome while red dots only represent the core interactome of the TBEV genome. Green triangles indicate viral proteins. The complete list of identified proteins is provided in Supplementary Table S2.

A total of 4531 proteins, the majority corresponding to non-specific signals, were detected by MS (Table S2). Those that specifically interact with the viral genome were discriminated according to the following criteria: at least 2 unique peptides for a given protein, at least 2-fold enrichment of peptides detected in the TBEV condition as compared with the NI condition and the p-values associated with peptide abundance for the TBEV condition set at a maximum of 0.05. This filtering qualified 822 proteins as specific interactors of the TBEV genome (Figure 1D, red + orange dots and Table S2), a set we termed its “extended interactome”. By lowering the statistical threshold to p-values ≤ 0.01, we then obtained 215 high-confidence interactors of the viral genome (Figure 1D, red dots and Table S2), a set we referred to as its “core interactome”.

As expected, multiple viral proteins were detected in this screen (Figure 1D, green triangles). These included several well-described viral RBP; namely, the RNA-dependent RNA polymerase/methyl transferase NS5, the protease/helicase NS3, the capsid C protein, which is part of the ribonucleocapsid core. NS2A was also detected, but its enrichment did not reach the p-value threshold (Table S2). NS2B, though not an RBP, was present in the highly enriched set of viral proteins, perhaps due to its interaction with NS3. The remaining non-structural proteins, which together form the viral replication complex, were also recovered, as was the structural protein prM. Our data cannot distinguish whether the viral proteins are detected as individual proteins or as part of the precursor viral polyprotein. The absence of the E protein from the data set, however, may suggest that the polyprotein was not retrieved during oligoprecipitation, and therefore that the viral proteins were recovered individually in their mature form.

### Characterization of the core and extended interactomes of the vRNA during TBEV infection

To characterize the core interactome, each interactor was manually annotated in reference to the existing literature as well as information from NCBI Gene (https://www.ncbi.nlm.nih.gov/gene), and Uniprot (https://www.uniprot.org/) databases. Of note, only 21% of the proteins have previously been cited in connection with binding to RNA. Based on data from several functional screens involving siRNA silencing, genome editing or overexpression, 79% of the discovered proteins had already been described as having a functional impact on the viral cycle, when all families of viruses were considered. However, the mechanisms by which the host proteins impinge upon viral replication were rarely characterized (Table S3).

In addition, Gene Ontology (GO) analysis was performed to retrieve enriched, statistically significant GO terms represented among the proteins in the core (Figure 2) and extended (Figure S1) interactomes. The most salient biological processes (BP), molecular functions (MF), and cellular components (CC), grouped in functional clusters, are shown.

**Figure 2.**
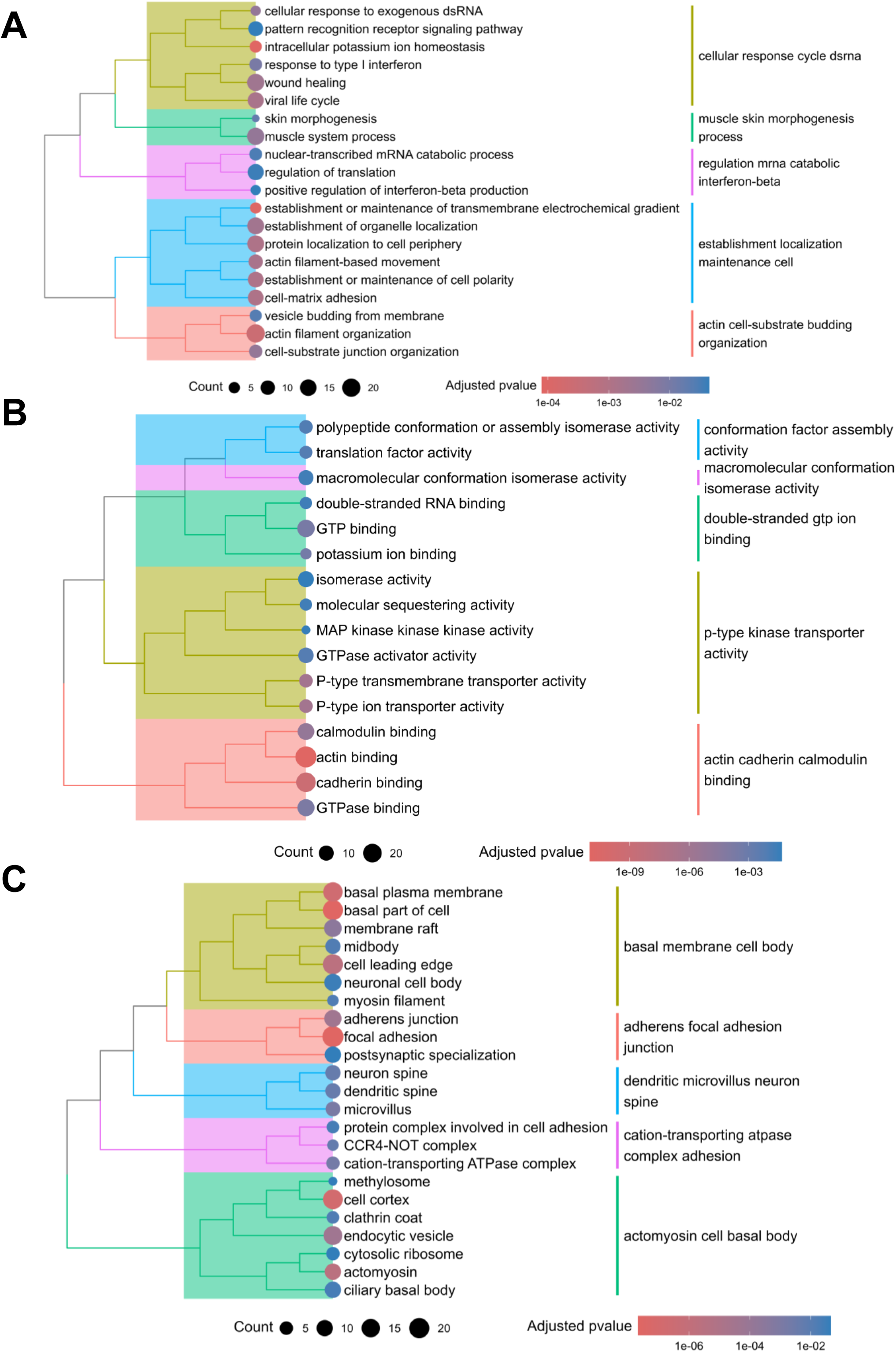
Biological processes, molecular functions, and cellular compartments engaged by the viral RNA genome during TBEV infection. Dendrograms depicting Gene Ontology enrichment for (A) biological processes, (B) molecular functions and (C) cellular compartments associated with the proteins in the core interactome of the TBEV genome. GO-terms are clustered based on semantic similarity. Circles size and color reflect the number of enriched proteins within each term and the associated adjusted p-value, respectively. Only terms associated with an adjusted p<0.05 were retrieved and displayed.

Among the over-represented BP, several are related to dsRNA sensing, pattern recognition receptor (PRR) signaling, and type I interferon (IFN) responses, reflecting the import of viral RNA as both trigger and target of innate antiviral immunity. The core interactome of TBEV RNA comprises multiple components of innate immune pathways. These notably include the retinoic acid inducible gene I (RIG-I)-like receptors (RLR) RIG-I (or DDX58) and Melanoma differentiation-associated gene 5 (MDA5) (or IFIH1), both key sensors of viral RNA, and the Signal transducer and activator of transcription 1 (STAT1), essential for induction of interferon-stimulated genes (ISG)^34,35^. Several ISG, such as ISG15, 2’-5’-Oligoadenylate Synthetase 3 (OAS3), Interferon Induced Protein With Tetratricopeptide Repeats 1 (IFIT1) and 2 (IFIT2) and MX Dynamin Like GTPase 1 (MX1) and 2 (MX2)^36^, are also found in the core interactome, as is Vesicle Associated Membrane Protein 8 (VAMP8), a SNARE protein recently implicated in the type I IFN response elicited by West Nile virus (WNV) infection. Moreover, in the extended interactome we note the presence of IFIT3 and Tripartite Motif Containing 25 (TRIM25), also involved in innate immune signaling^37^, as well as Ubiquitin Protein Ligase E3 Component N-Recognin 4 (UBR4), shown to be co-opted by DENV to inhibit IFN-I signaling via STAT2 degradation^38^.

Multiple over-represented BP are connected with mRNA catabolism and translational regulation, reflecting the role of host proteins, whether as pro- or antiviral factors in regulation of viral gene expression. Translation-associated factors are strongly represented in the interactome (Table S4-S5). Core interactors include the ribosomal proteins RPL23, RPL27, and RPS4X, and an additional twenty ribosomal proteins appear in the extended interactome. Three eukaryotic initiation factors—EIF4G2, EIF5, and EIF5B—are also part of the core set, with twelve more present in the extended one. These EIFs participate in the initiation phase of eukaryotic translation by stabilizing the assembly of ribosomal complexes. Components of the CCR4–NOT (CNOT) complex, which regulates translation and mRNA decay, are also represented, with CNOT1 and CNOT3 in the core interactome and CNOT9 as well in the extended set^39^. Moreover, three post-transcriptional modifiers, WDR4 TRNA N7-Guanosine Methyltransferase Non-Catalytic Subunit (WDR4), Elongator Acetyltransferase Complex Subunit 3 (ELP3) and DTD1, were also identified. High Density Lipoprotein Binding Protein (HDLBP) (Vigilin) and SERPINE1 MRNA Binding Protein 1 (SERBP1) (in the extended set) have been shown to link the vRNA of Dengue virus (DENV) to the host cell’s translation machinery, thereby promoting viral replication^17^. Finally, another interactor with antiviral activity, Exocyst Complex Component 1 (EXOC1), described as a transcriptional repressor of orthoflaviviruses through the sequestration of the elongation factor 1alpha (EF1alpha), was detected in the core interactome^40^.

Overrepresentation of BP connected with vesicular trafficking and cytoskeletal remodeling, as well as organelle localization, relates to interactions that engage vRNA during cytoskeleton-dependent trafficking or within endoplasmic reticulum (ER)-derived replication organelles. The vRNA interactome of TBEV comprises a large set of proteins with actin-, cadherin-, and calmodulin-binding activities, that mirror the dynamic cytoskeletal and adhesion processes required across the viral life cycle. For example, our screen uncovered the myosins MYO1B, MYO1C, MYO6, MYH10 and MYH14, motor proteins that move along actin filaments, producing the mechanical force required for intracellular transport, changes in cell shape or cell motility. Such motors are frequently co-opted by viruses, including TBEV, to support various steps of the viral life cycles^6,9,41^. We also detect the tropomyosins TPM1 and TPM3 in the core interactome, and TPM4 in the extended one, which are well-known stabilizers of actin filaments. Septin2, which is involved in DENV replication, is also part of the core interactome^42,43^. In parallel, several core interactome proteins map to vesicular trafficking pathways, including AP2A2, CAV1, EHD2, EPN1, FLOT2, and PICALM, which participate in caveolae- or clathrin-dependent endocytosis. We also detect ATL1 and ATL3, two ER-shaping atlastin proteins which induce ER membrane fusion, described as facilitating ZIKV replication^44^.

The cellular component analysis (Figures 2C and S1C) further links these functions within specific subcellular structures: enrichment of ER membranes, cytosolic ribonucleoprotein complexes, actin cytoskeleton, cell–substrate junctions, and vesicle-associated compartments reflect the formation of ER-derived replication organelles, the remodeling of cytoskeletal networks, and the engagement of trafficking pathways essential for viral replication and release. Of note, the core interactome includes several nuclear proteins involved in chromatin compaction and repair (H2BC5, H2BC9, XRCC4), protein import/export (NUP107, NUP188, NUP54), epigenetic modification (HDAC6) and transcriptional regulation (MED1, RTRAF). While replication of orthoflaviviruses occurs in the cytoplasm, we recently identified multiple nuclear host factors interacting with the viral RBP NS5.

Together, these GO signatures illuminate the host cell functions that TBEV exploits or endures in the course of its life cycle.

### Comparison of the TBEV RNA interactome with other orthoflaviviral interactomes uncovered by ChIRP-MS

The vRNA interactome of two mosquito-borne orthoflaviviruses (DENV and Zika virus (ZIKV)) with human cells has previously been resolved by ChIRP-MS methodology. One study addressed both DENV and ZIKV and generated a single data set, while the other only addressed ZIKV^12,13^. Both sets of proteins were compared with those from the present study (Figure 3A). Regarding the core interactome of TBEV, 15 interactors were shared with Ooi *et al.* (DENV & ZIKV), 3 with Zhang *et al.* (ZIKV) and 3 were common to all three. Unsurprisingly, the two data sets that relate to the RNA interactome of ZIKV display greater overlap (Figure 3A). Notably, none of these three data sets overlapped with that of Muto *et al.*, who addressed the interactome of a synthetic version of the 3’UTR for multiple TBEV strains in an *in vitro* context (not shown)^24^.

**Figure 3.**
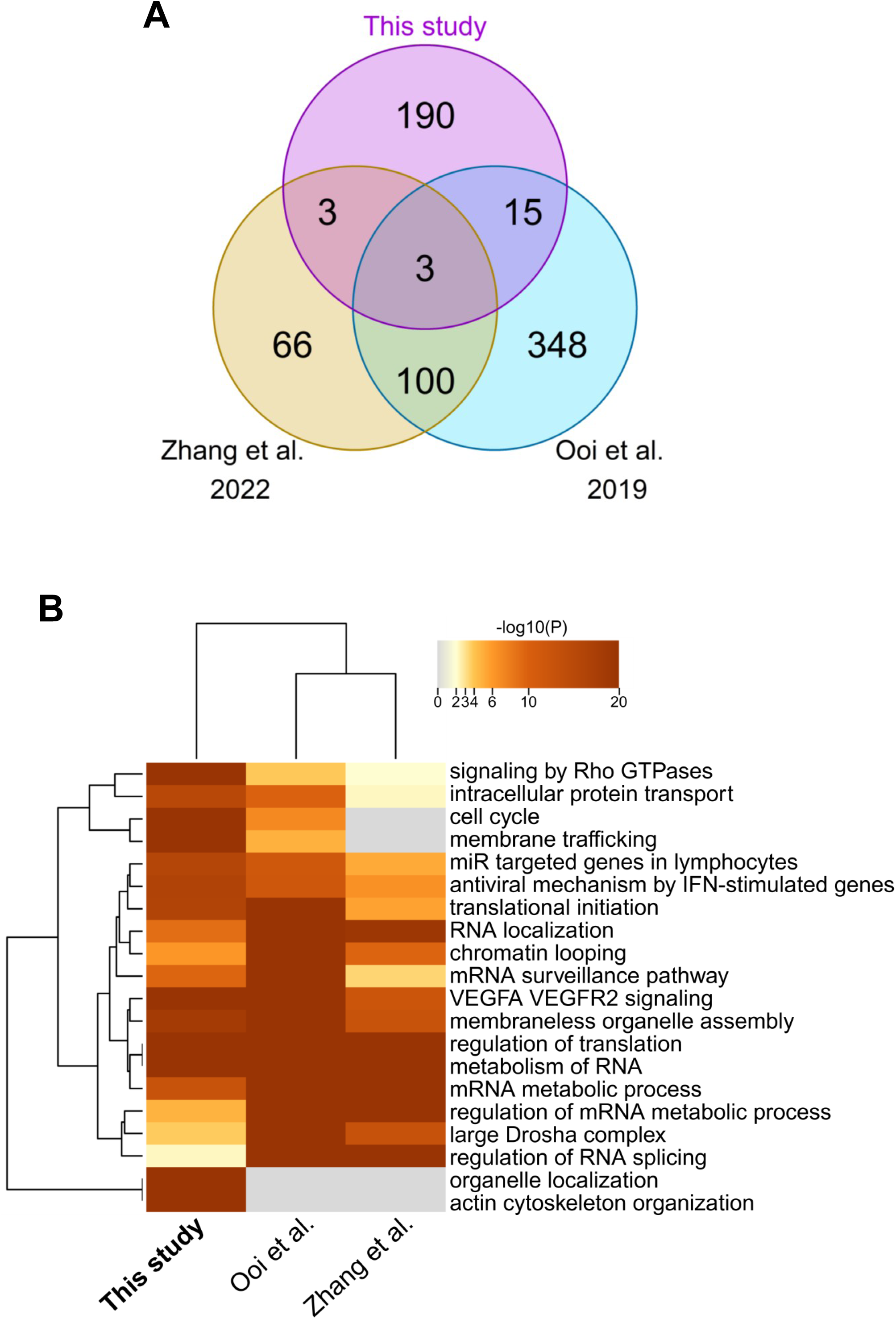
Comparison of the orthoflaviviral RNA interactome resolved in human cells by ChIRP-MS. (A) Venn-diagram showing the overlap between TBEV RNA interactors and those identified in two orthoflaviviral ChIRP-MS studies: Zika virus (Zhang *et al.* 2022) and dengue + Zika virus (Ooi *et al.* 2019). (B) Heatmap displaying the biological processes commonly enriched across the three orthoflaviviral interactomes.

Upon comparison of the BP of these three studies, we observed that GO terms related to RNA, transcription, vesicle trafficking and the type I IFN antiviral response were common to all three. Unexpectedly, however, GO terms related to actin cytoskeleton rearrangement and organelle localization were completely absent from the interactome of the two mosquito-borne orthoflaviviruses (Figure 3B).

### Functional analysis of the TBEV vRNA-host protein interactions identified in our screen

To address the implication of the human proteins identified in the present study in the TBEV life cycle, we suppressed their expression by RNA interference (RNAi) and examined the effect on viral replication. We focused on a representative set of 30 viral genome interactors, covering just over 10% of the core interactome.

Data from the GO term enrichment analysis and the manual annotation were integrated to select the 30 factors of interest, listed in Table 1, based on their enrichment level in our screen, their propensity for RNA binding, their involvement in cellular pathways or their implication in an orthoflaviviral life cycle.

**Table 1.** Summary of TBEV–*H. sapiens* RNA–protein interactors identified by ChIRP-MS and selected for functional analysis. Column A lists the gene names, Columns B and C provide the corresponding UniProt identifiers and UniProt protein descriptions. Column E reports the fold enrichment over non-infected control in the ChIRP-MS screen. Column F summarizes the roles of each interactor, and Column G indicates their RNA-binding protein (RBP) status. Column H compiles the supporting bibliography. Orange cells highlight factors enriched more than 100-fold in the screen; blue cells denote RBPs; and green cells mark interactors involved in cellular structural organization, one of the most strongly enriched GO term clusters.

| Gene | Uniprot ID | Name | Fold enrichment | Role | RBP | Reference |
| --- | --- | --- | --- | --- | --- | --- |
| AFDN | P55196 | Afadin | 3.80 | Nectin- and actin-filament-binding protein | NO | doi: 10.1016/s0378-1119(00)00316-4 |
| ASMTL | O95671 | Acetylserotonin O-Methyltransferase Like | 191.24 | Participates in nucleoside metabolism via its dTTP and UTP pyrophosphatase activity | NO | doi: 10.1016/j.chembiol.2013.09.011 |
| CAV1 | Q03135 | Caveolin-1 | 10.39 | Scaffolding protein within caveolar membranes | NO | doi: 10.3390/biology12111402 |
| CNOT1 | A5YKK6 | CCR4-NOT transcription complex subunit 1 | 4.25 | mRNA deadenylation | YES | doi: 10.1007/s13238-011-1092-4 |
| CNOT3 | O75175 | CCR4-NOT transcription complex subunit 3 | 2.82 | mRNA deadenylation | YES | doi: 10.1007/s13238-011-1092-4 |
| DDX58 | O95786 | Retinoic acid-inducible gene I | 3.46 | Cytosolic pattern recognition receptor (PRR) that can mediate induction of type-I IFN | YES | doi: 10.1016/B978-0-12-410524-9.00004-9 |
| DHX57 | Q6P158 | Putative ATP-dependent RNA helicase DHX57 | 6.65 | Probable ATP-binding RNA helicase | YES |  |
| DTD1 | Q8TEA8 | D-aminoacyl-tRNA deacylase 1 | 24.30 | Aminoacyl-tRNA editing enzyme | YES | doi: 10.1371/journal.pbio.1002465 |
| ELP3 | Q9H9T3 | Elongator complex protein 3 | 8.07 | tRNA acetylation | YES | doi: 10.1093/hmg/ddy043 |
| EXOC1 | Q9NV70 | Exocyst complex component 1 | 3.20 | Involved in the docking of exocytic vesicles | NO |  |
| GPATCH8 | Q9UKJ3 | G patch domain-containing protein 8 | 327.53 | Unknown | ? |  |
| HDLBP | Q00341 | high density lipoprotein binding protein / Vigilin | 3.99 | Plays a role in cholesterol metabolism | NO |  |
| HIBADH | P31937 | 3-hydroxyisobutyrate dehydrogenase | 407.48 | Role in the catabolism of L-valine | NO | doi: 10.1016/S0076-6879(00)24234-1 |
| IFIH1 | Q9BYX4 | Interferon-induced helicase C domain-containing protein 1 | 909.10 | Cytosolic pattern recognition receptor (PRR) that can mediate induction of type-I IFN | YES | doi: 10.1016/B978-0-12-410524-9.00004-9 |
| ILKAP | Q9H0C8 | Integrin-linked kinase-associated serine/threonine phosphatase 2C | 206.27 | Protein phosphatase that plays a role in regulation of the cell cycle | NO | doi: 10.1038/sj.onc.1207473 |
| MACF1 | Q9UPN3 | Microtubule-actin cross-linking factor 1 | 2.57 | F-actin-binding protein | NO | doi: 10.3389/fcell.2021.641727 |
| MX1 | P20591 | Interferon-induced GTP-binding protein Mx1 | 19.23 | Interferon-induced dynamin-like GTPase | NO | doi: 10.3390/vaccines11050930 |
| MX2 | P20592 | Interferon-induced GTP-binding protein Mx2 | 9208.79 | Interferon-induced dynamin-like GTPase | NO | doi: 10.3390/vaccines11050930 |
| MYH10 | P35580 | Myosin heavy chain 10 | 4.00 | Actin-dependent motor protein | NO |  |
| MYO1C | O00159 | Unconventional myosin-Ic | 8.23 | Actin-based motor molecule with ATPase activity | NO |  |
| NBAS | A2RRP1 | NBAS subunit of NRZ tethering complex | 121.50 | Involved in Golgi-to-endoplasmic reticulum (ER) retrograde transport | NO | doi: 10.1091/mbc.e08-11-1104 |
| NUMB | P49757 | Protein numb homolog | 944.26 | Regulates clathrin-mediated receptor endocytosis | NO | doi: 10.1111/j.1600-0854.2008.00790.x |
| PTDSS1 | P48651 | Phosphatidylserine synthase 1 | 8.37 | Catalyzes the formation of phosphatidylserine | NO | doi: 10.1042/BJ20081597 |
| RNF213 | Q63HN8 | E3 ubiquitin-protein ligase RNF213 | 3.52 | E3 ubiquitin ligase that can catalyze ubiquitination of both proteins and lipids | NO | doi: 10.1093/jb/mvae020 |
| RPS4X | P62701 | Small ribosomal subunit protein eS4, X isoform | 2.44 | Component of the small ribosomal subunit | NO | doi: 10.1038/nature12104 |
| RRBP1 | Q9P2E9 | Ribosome-binding protein 1 | 2.50 | Mediates interaction between the ribosome and the endoplasmic reticulum membrane | NO |  |
| SRGAP2 | O75044 | SLIT-ROBO Rho GTPase-activating protein 2 | 269.42 | Stimulates GTPase activity of Rac1, and plays a role in cortical neuron development | NO | doi: 10.1074/jbc.M110.153429 |
| STAU1 | O95793 | Double-stranded RNA-binding protein Staufen homolog 1 | 4.66 | Binds double-stranded RNA and tubulin | YES |  |
| WDR4 | P57081 | tRNA guanine-N(7)-methyltransferase non-catalytic subunit WDR4 | 193.95 | Part of a N7-methylguanosine transferase complex for tRNA, mRNA and microRNA | YES | doi: 10.1017/s1355838202024019 |
| ZYX | Q15942 | Zyxin | 10.02 | Adhesion plaque protein | YES | doi: 10.1016/j.ceb.2006.08.006 |

Among these proteins, ten displayed greater than 100-fold enrichment in our screen (Table 1 – orange cells), ten had been described as RBP (Table 1 – blue cells), and six interactors were involved in cellular structural organization (Table 1 – green cells), one of the most enriched GO term clusters. We also included interactors described to have an impact on orthoflavivirus life cycles, namely, HDLBP, which promotes DENV infection^17^, RRBP1 (found in the extended interactome), which is a proviral factor for several orthoflaviviruses, including the tick-borne orthoflavivirus Powassan^13^, EXOC1 known to repress DENV and WNV, although the capsid C protein antagonizes this antiviral activity^40^, MDA5 and RIG-I, two viral RNA sensors responsible for triggering the type I IFN response, and MX2 an ISG with antiviral activity for DENV and JEV^45^. The ISG MX1, though not known to display antiviral activity against orthoflavivirus, was added for comparison with MX2. We also knocked down several genes whose cognate proteins were not retrieved in our screen. Three RBP, NCL, NONO, and YBX1, were included owing to their proviral activity against several orthoflaviviruses^46–48^. Finally, TMEM41B, which is required for the multiplication of many orthoflaviviruses, including TBEV, was included as a control for proviral activity^49^.

Expression of the proteins of interest was knocked down in A549 cells 48h prior to infection with TBEV, and viral replication was assessed 48h later by quantification of infectious particles released into the supernatant (Figure 4A). Transfection with irrelevant (non-targeting) siRNA (siNT) was used as a reference. Cell viability after each treatment was verified by ATP measurement. Proteins whose silencing induced more than 25% mortality, such as MACF1 and NUMB, were excluded from further experimentation (Figure S2A). The silencing efficiency of our factors was also verified by RT-qPCR in comparison to that of siNT (Figure S2B). Proteins whose silencing efficiency was less than 50%, such as AFDN, HIBADH, MYO1C and NUMB, were also excluded from further consideration.

**Figure 4.**
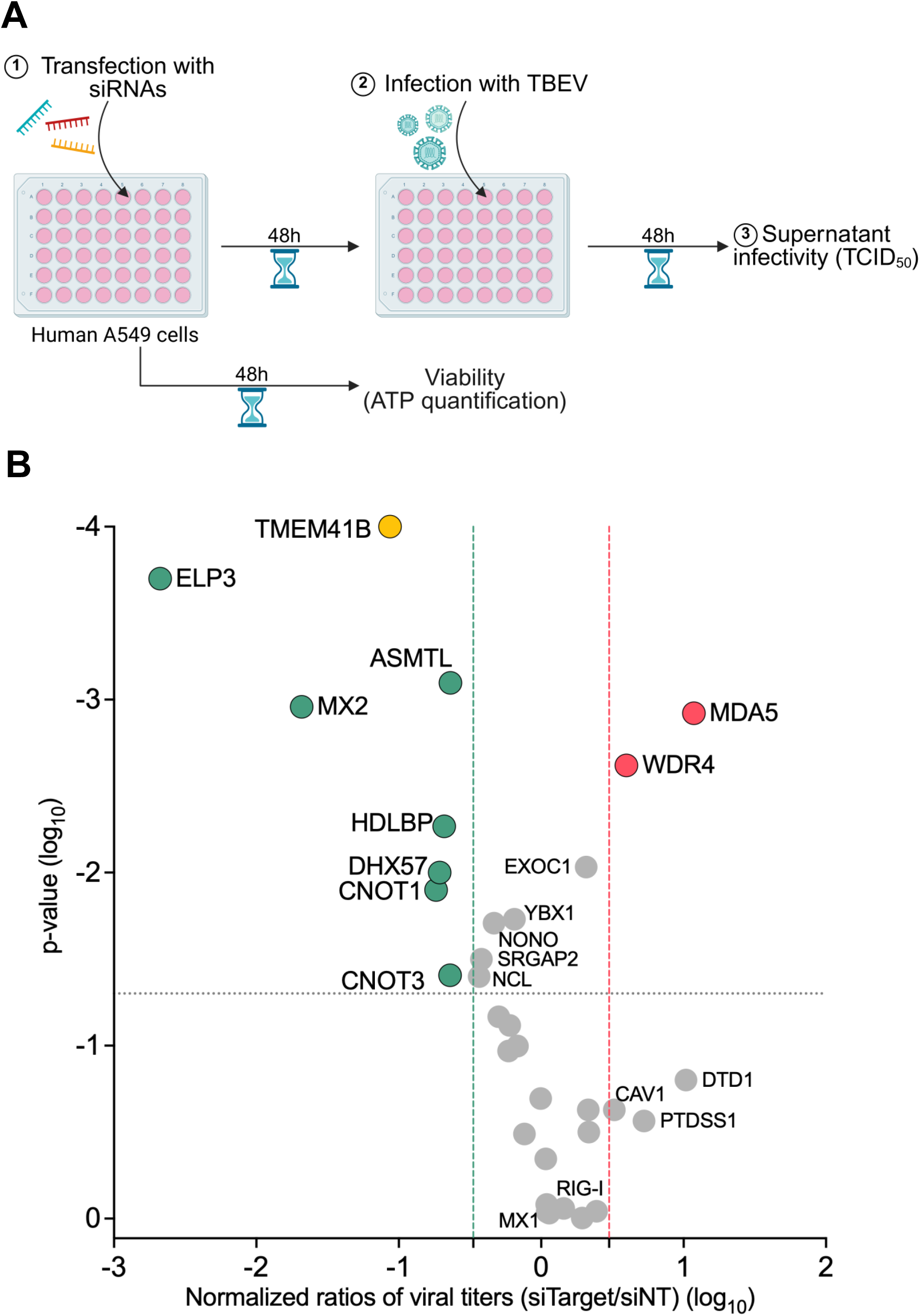
Functional analysis of the TBEV RNA interactome reveals novel dependency and restriction hosts factors. (A) Schematic overview of the RNAi gene-silencing workflow (B) Scatter plot showing the fold change in infectious viral particle release by siRNA-silenced and TBEV-infected A549 cells, against statistical significance. Fold change is calculated as the mean viral titer in siTarget-transfected cells relative to controls siNT. Host factors are considered to significantly impact viral fitness if the titer ratio increases or decreases by at least 3-fold, with an adjusted p-value<0.05. Green, red, grey, and yellow dots represent proviral, antiviral, non-effective, and control proteins, respectively. Data represent means of 3 independent experiments, each performed in technical triplicates. Statistical significance was assessed using the non-parametric Kruskal–Wallis test, followed by Dunn’s multiple-comparison *post hoc* test with Benjamini–Krieger–Yekutieli two-stage step-up correction for multiple testing.

Infectious titers were normalized to the titers obtained after treatment with siNT and plotted against their corresponding p-values (Figure 4B). A host factor was classified as a hit when silencing induced at least a threefold change in normalized infectivity, with a p-value <0.05. Host factors whose silencing decreased titers were designated as dependency factors, while those whose silencing increased titers were designated as restriction factors.

Of the 30 factors tested, ten had a statistically significant effect (p<0.05) on TBEV replication (Figure 4B). Of the seven interactors that displayed proviral activity, ELP3 was the strongest, displaying a 474-fold decrease in infectivity when knocked down. Strikingly, MX2, a well-known antiviral protein, showed a strong proviral effect, reducing TBEV supernatant infectivity by 48-fold. As expected, in keeping with observations for the Hepatitis C virus (HCV)^50^, CNOT1 and CNOT3 enhanced TBEV replication, as did HDLBP. Of note, DHX57 and ASTML, which had never been described as affecting a viral cycle, facilitated production of infectious viral particles. While NONO did not reach the cut-off that we arbitrarily set for proviral activity, its knockdown reduced TBEV infectivity by more than 2-fold. In contrast, RRPB1 and YBX1 had little to no effect on TBEV infection. The proviral control TMEM41B acted as expected, its loss of expression reducing TBEV infectivity by 11.6-fold.

MDA5, an immune sensor of multiple RNA viruses, had a strong antiviral effect on TBEV replication, restricting TBEV infectious release by 11.8-fold, whereas with only a 1.4-fold increase in TBEV infectivity when silenced, RIG-I did not restrict viral replication under our experimental conditions. Of note, WDR4, a nuclear protein specialized in RNA modification as part of the tRNA N7-guanosine methyltransferase complex, is the only tested interactor that significantly restricted TBEV (by 4-fold). We note that knockdown of DTD1 and PDTSS1 increased infectivity by 4- and 5.3-fold, respectively, without attaining statistical significance. EXOC1 did not impair TBEV replication, despite its restriction of mosquito-borne viruses. Whether TBEV in particular or tick-borne orthoflaviviruses in general are insensitive to EXOC1, or fully antagonize EXOC1 restriction, remains to be determined.

### Host restriction and dependency activity of TBEV interactors against other arboviruses

Given the results of this functional analysis and the puzzling phenotype of some interactors, such as MX2 or RIG-I, in TBEV infection, we wished to determine whether the restriction and dependency factors for TBEV-specific or conserved across different positive-stranded RNA viruses. Of note, in a previous study, we demonstrated that tick-borne but not mosquito-borne orthoflaviviruses antagonize the type I IFN pathway by means of an interaction between NS5 and mammalian Tyk2^7^.

To investigate the breadth of activity for pro- and antiviral factors (Figure 5), we tested the susceptibility of three single-stranded positive RNA viruses: Louping Ill Virus (LIV), JEV, and CHIKV. LIV is a tick-borne orthoflavivirus that is phylogenetically close to TBEV, displaying up to 95% protein sequence identity and also transmitted in Europe by *Ixodes ricinus*^51^. JEV is also an encephalitic orthoflavivirus, but that is transmitted by *Aedes* and *Culex* mosquitoes. Finally, CHIKV belongs to a different family, the *Togaviridae*, making it phylogenetically distinct from TBEV, and is transmitted by *Aedes* mosquitoes.

**Figure 5.**
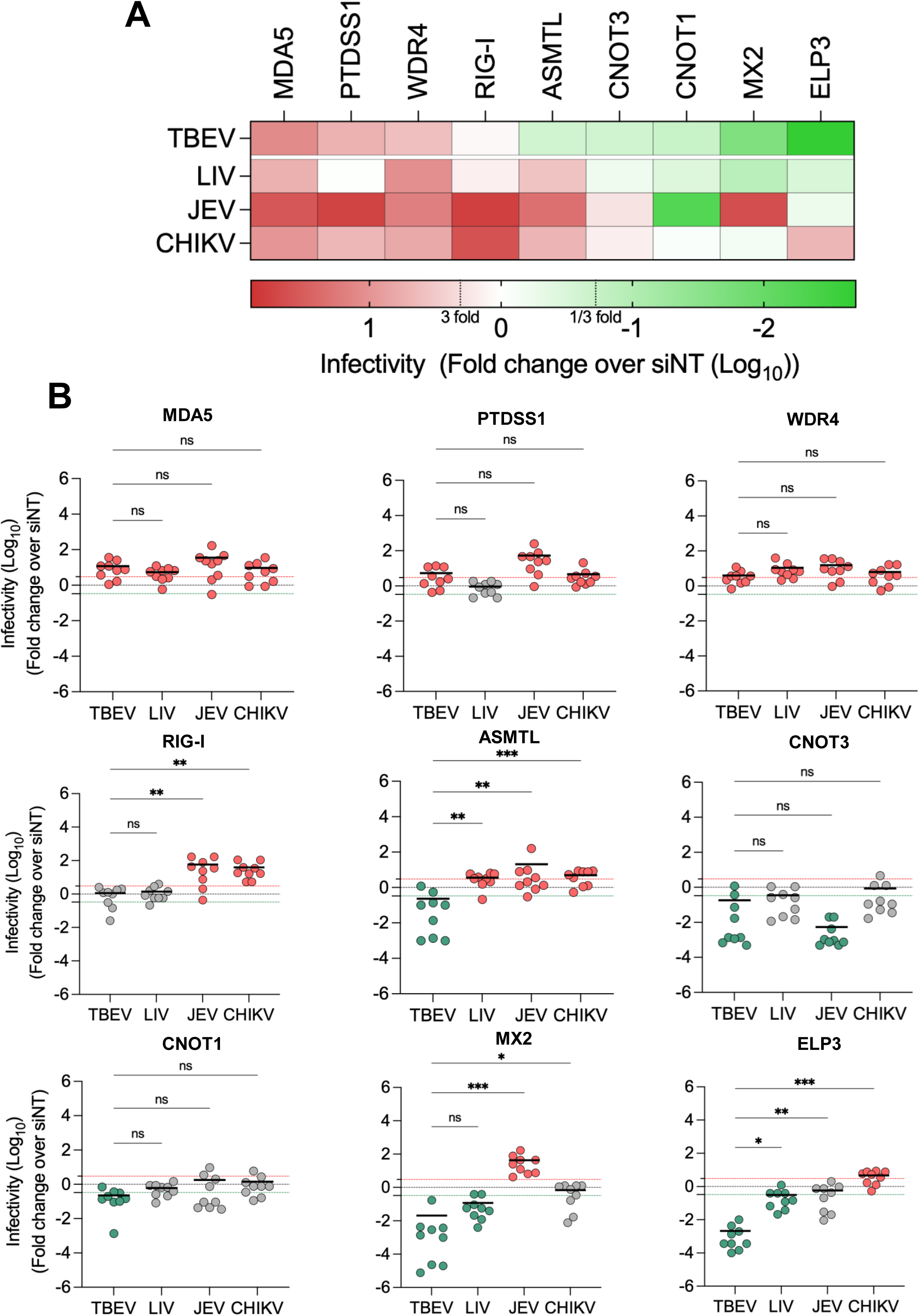
Host dependency and restriction factors of TBEV infection are shared among other arboviruses. (A) Heatmap summarizing the impact of RNAi-mediated silencing of each tested host factor on the replication cycles of LIV, JEV, and CHIKV relative to TBEV. Green and red tiles indicate proviral and antiviral factors, respectively. (B) Dot plots showing the individual effects of each tested host factor on viral replication. Green, red, and grey dots correspond to proviral, antiviral, and non-effective proteins, respectively. Statistical significance was assessed using the non-parametric Kruskal–Wallis test, followed by Dunn’s multiple-comparison *post hoc* test with Benjamini–Krieger–Yekutieli two-stage step-up correction for multiple testing (ns: non-significant; *p < 0.01, **p < 0.001, ***p < 0.0001).

The impact of silencing on replication of LIV, JEV or CHIKV was determined in A549 cells (Figure 5A), as described before for TBEV. As a general observation, the impact of the factors varied according to phylogenetic distance from TBEV: the more distant the viruses were phylogenetically, the less conserved were the phenotypes. (Figure 5A).

Among the host factors that restricted TBEV, WDR4 and MDA5 maintained their restriction against all three viruses (Figure 5A-B and S3). While the antiviral activity of MDA5 was expected to be broad, WDR4 is a newly discovered restriction factor with a seemingly wide spectrum of activity.

PTDSS1 restricted all viruses except LIV, with a major impact on JEV, for which its knockdown increased infectivity by 53-fold. Whereas RIG-I has been described to restrict TBEV in previous studies, under our experimental conditions it did not affect the replication of tick-borne viruses, though strongly restricting JEV (58-fold) and CHIKV (39-fold) (Figure 5A-B and S3).

ASMTL, which is proviral for TBEV, appears to be antiviral against the other three pathogens. TBEV may have evolved to evade the antiviral activity of ASMTL and subvert the interaction. CNOT3 only affected TBEV replication, while CNOT1 had no effect on LIV or CHIKV but strongly favored the replication of JEV (by 185-fold). Of note, the proviral phenotype of MX2 was conserved between LIV and TBEV, the two tick-borne orthoflaviviruses. In contrast, MX2 behaved, as expected, as a potent (43-fold) inhibitor of JEV replication, and had no impact on CHIKV. Finally, ELP3 was as proviral factor for both tick-borne orthoflaviviruses, while it restricted CHIKV and appeared to be neutral for JEV. (Figure 5A-B and S3). Collectively, these data suggest that in the course of their divergent evolution, tick- and mosquito-borne orthoflaviviruses have acquired the capacity to co-opt distinct host factors.

### WDR4, a new restriction factor of TBEV

Among the factors identified as influencing the replication of all tested arboviruses (Figure 5), WDR4, whose relation to viral infection is completely undescribed, was selected for further study. To confirm the restriction activity observed in the silencing experiments (Figures 4 and 5), we generated WDR4-knockout A549 cells (WDR4-KO A549) using CRISPR/Cas9 technology (Figure 6A) and performed TBEV infection assays (Figure 6B). 48 hours post TBEV infection, the infectious titer of the supernatant of WDR4-KO A549 cells was 97 times greater than that of the parental A549 WT cells. Moreover, stable re-expression of WDR4 via lentiviral transduction in WDR4-KO A549 cells restored infection to a level comparable to that of WT A549 cells, whereas expression of GFP as a control had only a modest effect on viral infectivity (Figure 6A-B). Altogether these results confirm the potential role of WDR4 as a restriction factor on TBEV infection.

**Figure 6.**
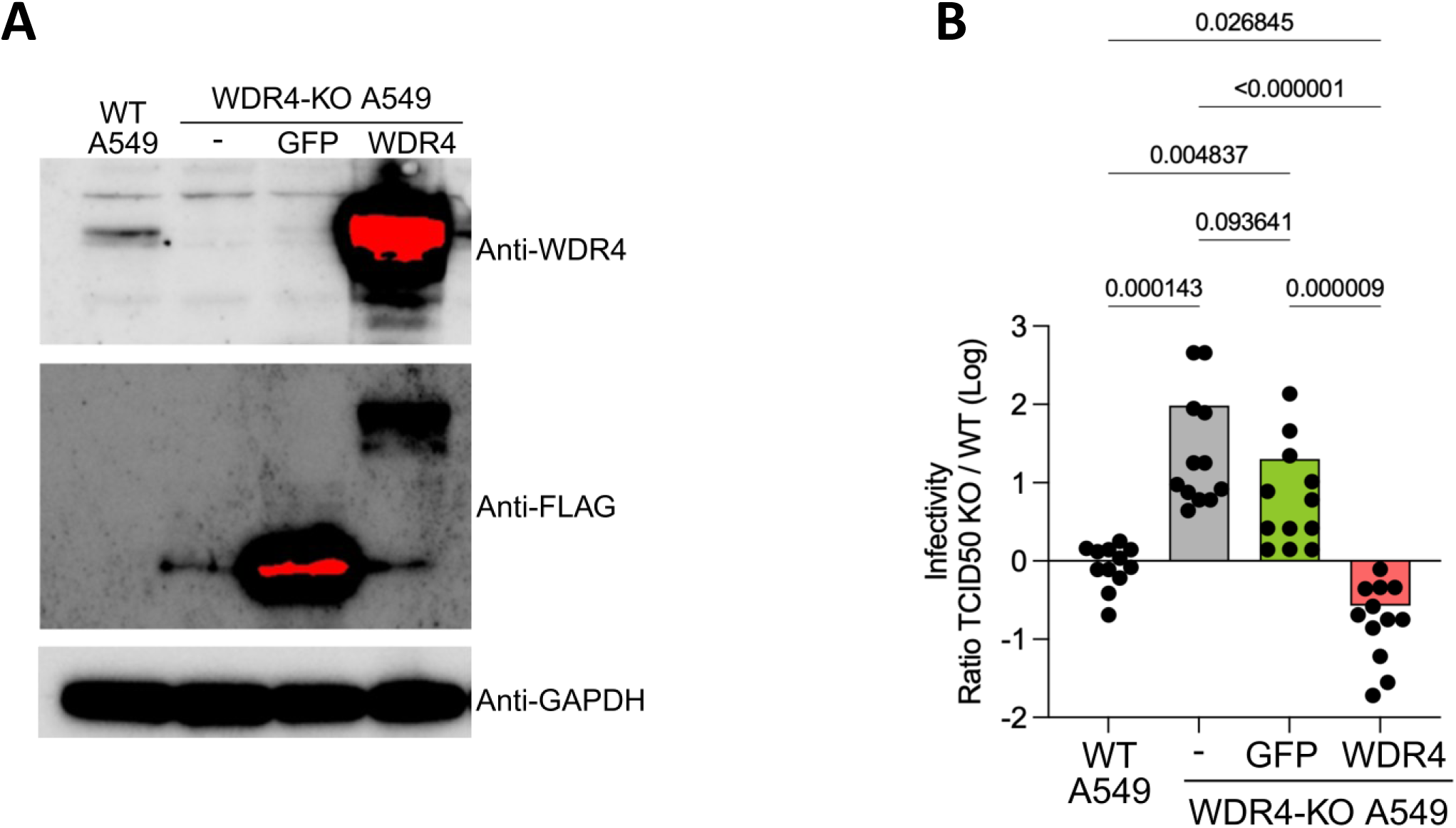
WDR4 restricts TBEV replication. (A) Expression levels of WDR4, GFP and GAPDH in the four indicated cell lines. (B) Comparison of TBEV infection levels in A549 wild type cells, WDR4 knockout A549 cells (WDR4-KO) and WDR4-KO cells stably expressing either GFP or WDR4. Data represent means of 4 independent experiments, each performed in technical triplicates. Statistical significance was assessed using the non-parametric Kruskal–Wallis test, followed by Dunn’s multiple-comparison *post hoc* test with Benjamini–Krieger–Yekutieli two-stage step-up correction for multiple testing (ns: non-significant; *p < 0.01, **p < 0.001, ***p < 0.0001).

## Discussion

The RNA genome of TBEV, as both the substrate for genome replication and translation and a designated target for innate antiviral immunity, is a nexus for interactions with the proteome of host cells. We have systematically mapped these interactions in infected human cells, thereby identifying a core interactome of 215 human host factors, which we functionally annotated. For selected RBP, we addressed their impact on TBEV infection in gene knockdown experiments, thereby identifying a set of restriction and dependency factors, whose impact was in some instances unexpected. These included sensors and effectors of innate immune pathways, as well as epitranscriptomic modifiers. Among the former, the ISG MX2, a dynamin-like GTPase displayed an unexpected proviral activity. Among the latter, WDR4, the non-catalytic component of the METTL1-WDR4 methyltransferase complex, emerged as a restriction factor with broad-spectrum activity against arboviruses belonging to multiple families of positive-strand RNA viruses.

To identify the proteins associated with the positive-strand TBEV RNA, we used ChIRP-MS methodology, which is based upon formaldehyde cross-linking followed by antisense oligonucleotide capture and mass spectrometry^25^. Of note, unlike UV cross-linking, formaldehyde fixation leads to the capture of not only direct RNA-binding proteins, but also proteins engaged in RNA-containing complexes, thereby offering a broader view of the viral RNA interactome. Moreover, as positive-strand RNA of orthoflaviviruses is both genome and mRNA, we expected to retrieve host proteins that encountered functionally and spatially distinct forms of vRNA.

### Functional annotation

Overrepresentation analysis of GO terms associated with the vRNA interactome of TBEV returned a set of biological processes, molecular functions and cellular components that are coherent with current understanding of the cellular biology of TBEN infection. The enrichment of processes such as actin filament organization, actin-based movement, cell–matrix adhesion, and cell polarity reflects the extensive interactions with the cytoskeleton required for viral entry, intracellular transport, replication in viral factories, and egress. The presence of focal adhesions, adherens junctions, membrane rafts, and actomyosin structures among the CC GO terms further reinforces the central role of adhesion and cytoskeletal networks in viral infection.

The importance of membrane trafficking pathways is further highlighted by the overrepresentation of GO terms associated with vesicle budding from membrane, establishment of organelle localization, and protein localization to the cell periphery. Orthoflaviviruses enter cells through clathrin- or caveolae-dependent endocytosis and then remodel ER membranes to generate replication organelles, a process that depends upon both vesicle dynamics and cytoskeleton-dependent organelle positioning^44,52–56^. The identification of clathrin coats, endocytic vesicles, and membrane microdomains among the CC terms aligns with this requirement.

Enriched BP GO terms also included translation regulation, mRNA catabolism, ribosomal components, and the CNOT complex, which evidence the immersive relationship with host protein and RNA metabolism during infection. Indeed, orthoflaviviral mRNA is translated at the ER membrane by hijacked ribosomal machinery^57,58^. Orthoflaviviruses also modulate host RNA decay pathways to enhance viral RNA stability. In this regard, CNOT complex components have been shown to promote the replication of HCV, a member of the *Flaviviridae* family. These findings align with broader evidence that orthoflaviviruses like TBEV exploit host translation machinery while regulating RNA quality-control systems to support replication.

Finally, the enrichment of GO terms related to pattern recognition receptor signaling, response to type I IFN, dsRNA sensing, and IFN-β induction reflects interaction with host innate immune pathways, which are strongly activated by viral infection. Positive-strand RNA orthoflaviviruses, such as TBEV, generate cytosolic dsRNA intermediates that trigger RLRs and downstream signaling^59,60^. TBEV delays and dampens the type I IFN response, notably by interfering with Tyk2 and STAT1/STAT2 signaling and ISG induction^7,61–64^. Of note, recent work shows that orthoflaviviruses also manipulate mitochondrial processes so as to modulate antiviral signaling, underscoring the interplay between viral replication and innate immunity^65^.

While a variety of host factors were identified in our screen, few of these were found in equivalent screens for other orthoflaviviruses. Nevertheless, comparison of overrepresented GO terms across the data sets revealed a globally conserved functional signature. Presumably tick-and mosquito-borne orthoflaviviruses engage the same pathways, but not necessarily via interactions with identical host proteins. Curiously, functional annotation of the interactome of two mosquito-borne orthoflaviviruses, though also resolved by ChIRP-MS, did not return overrepresented GO terms related to cytoskeleton and organelle localization. Presuming the requirement for cytoskeletal components at viral entry, viral translocation and egress to be nonnegotiable for all orthoflaviviruses, their absence from the data sets of mosquito-borne viruses is likely to reflect technical or analytical differences affecting the oligoprecipitation of host cell proteins or their identification as enriched proteins, respectively.

### Functional analysis

Functional interrogation of 30 selected interactors by RNA interference revealed multiple host factors that either restrict or promote TBEV replication. Notably, some of these factors have not previously been implicated in viral infection, highlighting the value of RNA-centric approaches to uncover noncanonical regulators of the viral life cycle. Of note, some host factors exhibit activities that differ from those reported for other viral or cellular systems, suggesting that their functional role may be highly context-dependent and influenced by virus-specific replication strategies or cellular environments. Our comparative analyses across additional orthoflaviviruses and alphaviruses, including both tick- and mosquito-borne viruses, revealed that restriction factors tended to display conserved activity across viruses, whereas proviral factors were more virus-specific, being restricted to orthoflaviviruses and, in some cases, to tick-borne viruses. We speculate that a greater conservation of restriction factors might be related to cellular ergonomy; that is, antiviral processes have been selected to counter the set of multiple and rapidly evolving pathogens found in their environment, while the multiplicity of dependency factors reflects the distinctive replication strategies of different viral families.

Among the host factors that displayed a marked effect on viral replication, several belonged to innate immune pathways, whether as sentinels or effectors. RLR are required for induction of type I IFN upon infection of various orthoflaviviruses, including TBEV^60,66,67^. The canonical RLR, RIG-I and MDA5, play non-redundant roles in the type I IFN response to orthoflaviviral infection^67^, notably through recognition of distinct structural elements in viral RNA. While both RIG-I and MDA5 are described to recognize double-stranded RNA, the former senses tri- and diphosphates exposed at the 5’ extremity of uncapped viral RNA, while the latter senses longer RNA species.

In our study, both MDA5 and RIG-I were significantly enriched in the set of host proteins associated with TBEV RNA. Nevertheless, knockdown of MDA5 but not RIG-I markedly increased the production of infectious viral particles for TBEV and the closely related tick-borne orthoflavivirus LIV, while knockdown of either RLR significantly enhanced amplification of the mosquito-borne virus JEV. Although in a previous study RIG-I rather than MDA5 was necessary for induction of IFN-β RNA during TBEV infection^60^, our results strongly suggest that MDA5 is also able to sense RNA of TBEV and the related tick-borne orthoflavivirus LIV and to trigger an antiviral signaling cascade. As the two studies differ in many qualitative and quantitative details—different cell lines, different quantities of virus, different timepoints, different biological criteria—direct comparison is precluded. Further studies will thus be needed to understand the relative contribution of MDA5 and RIG-I to sensing TBEV RNA and restricting TBEV replication.

Human MX2, which belongs to the dynamin superfamily of large GTPases, has been described as a restriction factor for multiple viruses, including HIV, HCV and the orthoflaviviruses JEV and DENV^68^. Depending on the virus, however, different mechanisms have been proposed. By binding to the HIV-1 capsid protein, MX2 reportedly sequesters the viral capsid within nucleoporin-containing cytoplasmic biomolecular condensates^69^. Restriction of HCV has been found to depend upon cyclophilin A (CypA); in particular, when bound to the NS5A protein of HCV, MX2 impeded binding between NS5A and CypA and subsequent translocation of NS5A to the endoplasmic reticulum^45^. Beyond HCV, the same authors report that MX2 exerts antiviral activity on two other members of the Flaviviridae family; that is, JEV and DENV, possibly through binding to the NS5 protein^45^.

MX2 was highly enriched in the set of proteins binding to TBEV RNA. Though not described as having RNA-binding capacity, MX2 may have been recovered with TBEV RNA by virtue of its interaction with an RNA-binding protein or RNA-binding protein complex. Of note, though MX2 has been described as interacting with the NS5 protein of the mosquito-borne orthoflaviviruses JEV and DENV, interaction with the NS5 protein of TBEV and LIV was not observed in our yeast 2-hybrid screen and affinity pulldown experiments^6,7^.

MX2 displayed unexpected proviral activity for the tick-borne orthoflaviviruses TBEV and LIV, despite restriction of the mosquito-borne orthoflavivirus JEV. The molecular mechanisms by which MX2 promoted viral replication are enigmatic. Though suggested that MX2 restriction of viruses belonging to the *Flaviviridae* family is related to dependency on CypA^45^, CypA has been described to be an important host factor for TBEV^70^. Otherwise, in multiple instances viruses have retasked canonical antiviral effectors to their own advantage^71^, and the proviral activity of the MX2 protein may possibly be an additional example. Finally, the proviral activity of MX2 may be related to a homeostatic function of the MX2 protein that has been hijacked by the virus. In this regard, though strongly induced by type I IFN, MX2 is constitutively expressed in some cells, and has been shown to play a role in mitochondrial integrity^72^.

Elucidating the mechanisms by which MX2 promotes replication of TBEV, while inhibiting that of JEV, will require extensive investigation that lies beyond the scope of the current study.

A second category of host factors displaying a pronounced effect on viral replication were post-transcriptional modifiers. Two such proteins, WDR4 and ELP3, are highly conserved eucaryotic proteins that play essential roles in the epitranscriptomic modification of host cell RNA. As part of the Elongator complex, ELP3 catalyzes acetylation of transfer RNA (tRNA) at the U34 position, thereby facilitating the wobble interactions required for decoding of mRNA during translation^73^. WDR4 is a non-catalytic component of the METTL1-WDR4 methyltransferase complex, which is responsible for the formation of 7-methyl guanosine (m7G) in several RNA species, notably in a subset of tRNA at position 46. Through its influence on steady-state tRNA levels, the heterodimeric m7G writer is an important regulator of cell growth.

It is as yet unclear whether the impact of ELP3 and WDR4 on TBEV replication is related to epitranscriptomic modification, but if so, could conceivably concern the epitranscriptome of either cellular or viral RNA. Owing to their role in sustaining efficient translation, diminution of ELP3 or WDR4 by gene knockdown might be expected to suppress viral production, as was the case for ELP3 but not WDR4, whose knockdown enhanced viral production. Nevertheless, only a subset of tRNA is methylated by the m7G writer, such that knockdown of WDR4 would selectively diminish the translation of mRNA species that are enriched in specific codons, rather than globally affecting translation^74^.

Though neither ELP3 nor WDR4 has been described to modify viral RNA, their enrichment in the viral RNA-binding subset of host proteins incites speculation. Flaviviral RNA is known to be bear multiple epigenetic marks, whether generated by the viral methyltransferase or by the host epigenetic machinery^75^, but to the best of our knowledge acetylation of uridine has not been documented. As regards WDR4, the catalytic component of the m7G writer, METTL1, was recovered along with viral RNA, but was not enriched in a statistically significant manner. Whether the antiviral effect of WDR4 is directly related to METTL1-WDR4-mediated N7-guanosine methylation of viral RNA will require further investigation.

Finally, the impact of ELP3 and WDR4 on viral production might be entirely unrelated to their role in translation. In this regard, ELP3 has recently been found to be essential for induction of the type I IFN response in human macrophages, by influencing not only PRR-mediated induction of IFN-β but also subsequent IFN-β-mediated activation of STAT1^76^.

### Conclusion

By resolving the interactome of TBEV RNA and the human proteome, we provide the first description of an RNA interactome for a tick-borne orthoflavivirus. The functional signature of vRNA-associated RBP is highly consistent with current models of orthoflavivirus–host interaction. Based on the subset of host RBP whose importance in viral infection was experimentally addressed, a large proportion possess pro- or antiviral activity. Among the various impactful host factors, the epitranscriptomic modifier WDR4 and immune effector MX2, novel restriction and dependency factors, especially warrant further investigation.

All together, these findings provide a global view of the TBEV RNA interactome and identify host pathways that are differentially exploited or antagonized during infection, thereby expanding our understanding of tick-borne orthoflavivirus biology.

## Figure legends

**Figure S1 Biological processes, molecular functions, and cellular compartments engaged by the viral RNA genome during TBEV infection**

Dendrograms depicting Gene Ontology enrichment for (A) biological processes, (B) molecular functions and (C) cellular compartments associated with the proteins in the extended interactome of the TBEV genome. GO-terms are clustered based on semantic similarity. Circles size and color reflect the number of enriched proteins within each term and the associated adjusted p-value, respectively. Only terms associated to an adjusted p<0.05 were retrieved and displayed.

**Figure S2 Viability and silencing efficiency of siRNA-treated cells.**

(A) Cell viability and (B) gene-silencing efficiency were assessed in A549 cells transfected with siRNAs targeting the indicated host factors. All values are expressed relative to the siNT treated cells. Data represent the mean ± SEM of three biological replicates.

**Figure S3 Functional analysis of impactful host factors for TBEV across diverse arboviruses**

Scatter plot showing the fold change in infectious viral particle release by siRNA silenced and (A) LIV-, (B) JEV- and (C) CHIKV-infected A549 cells, against statistical significance. Fold change is calculated as the mean viral titer in siTarget-transfected cells relative to siNT controls. Host factors are considered to significantly impact viral fitness if the titer ratio increases or decreases by at least 3-fold, with an adjusted p-value<0.05. Green, red, grey, and yellow dots represent proviral, antiviral, non-effective, and control proteins, respectively. Data represent means of 3 independent experiments, each performed in technical triplicates. Statistical significance was assessed using the non-parametric Kruskal–Wallis test, followed by Dunn’s multiple-comparison *post hoc* test with Benjamini–Krieger–Yekutieli two-stage step-up correction for multiple testing.

## Acknowledgments

This study was made possible by a grant from the French national research agency (Project N° ANR-24-CE35-7666-01). AG received doctoral funding from the Laboratoire d’excellence “Integrative Biology of Emerging Infectious Diseases”, in the framework of the “Investissements d’Avenir” program, as managed by the French national research agency (Reference N° ANR-10-LABX-62-01). The Ph.D. of ML is co-financed by the Domaine d’Intérêt Majeur 1Health of Ile de France and the Alfort Veterinary School.

