## Supplemental Figures 1-3 for "An RNA-centric interactomics screen identifies novel proviral and antiviral genome binding interactors for tick-borne encephalitis virus"

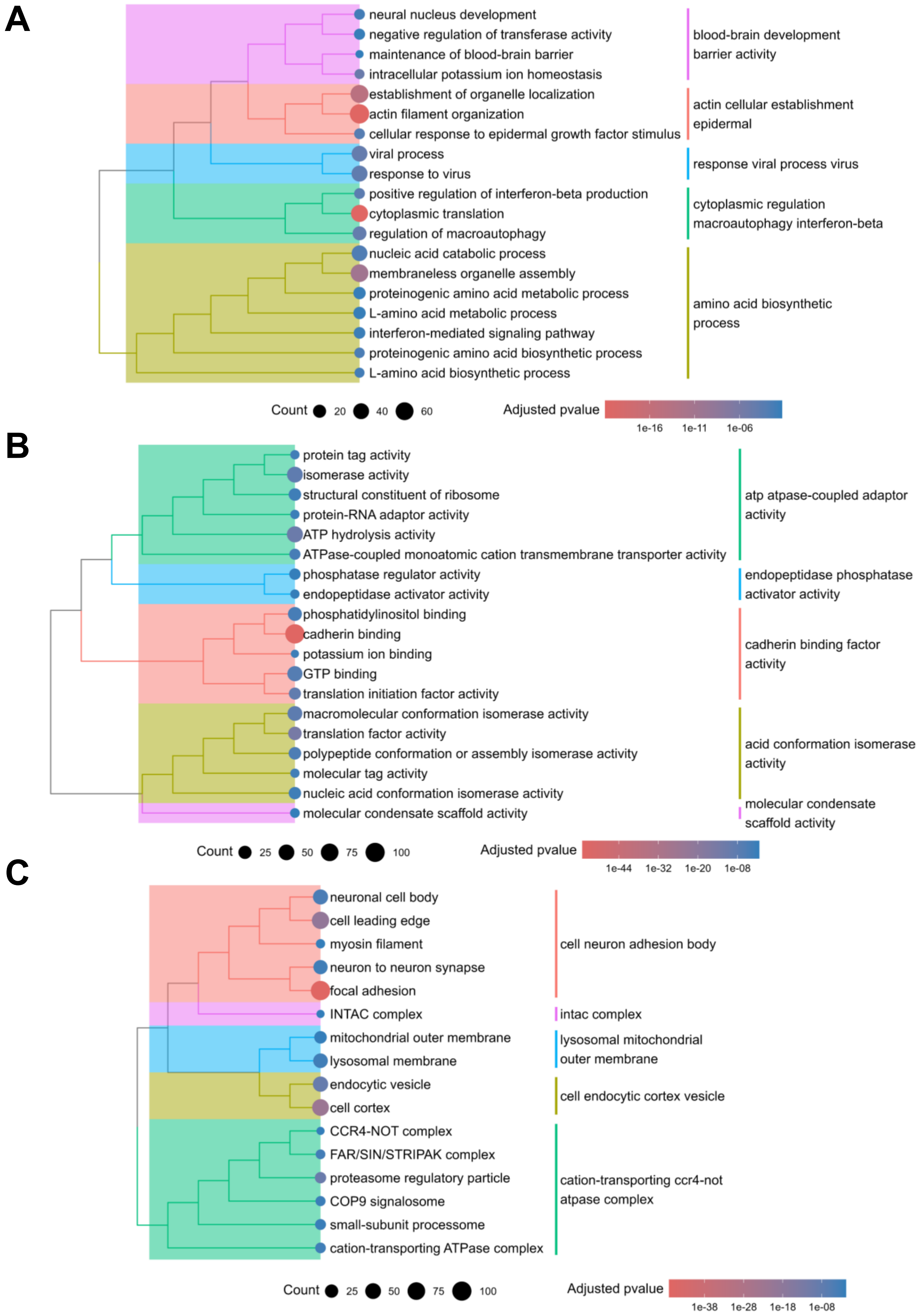


**Figure S1 - Biological processes, molecular functions, and cellular compartments engaged by the viral RNA genome during TBEV infection**

**A**

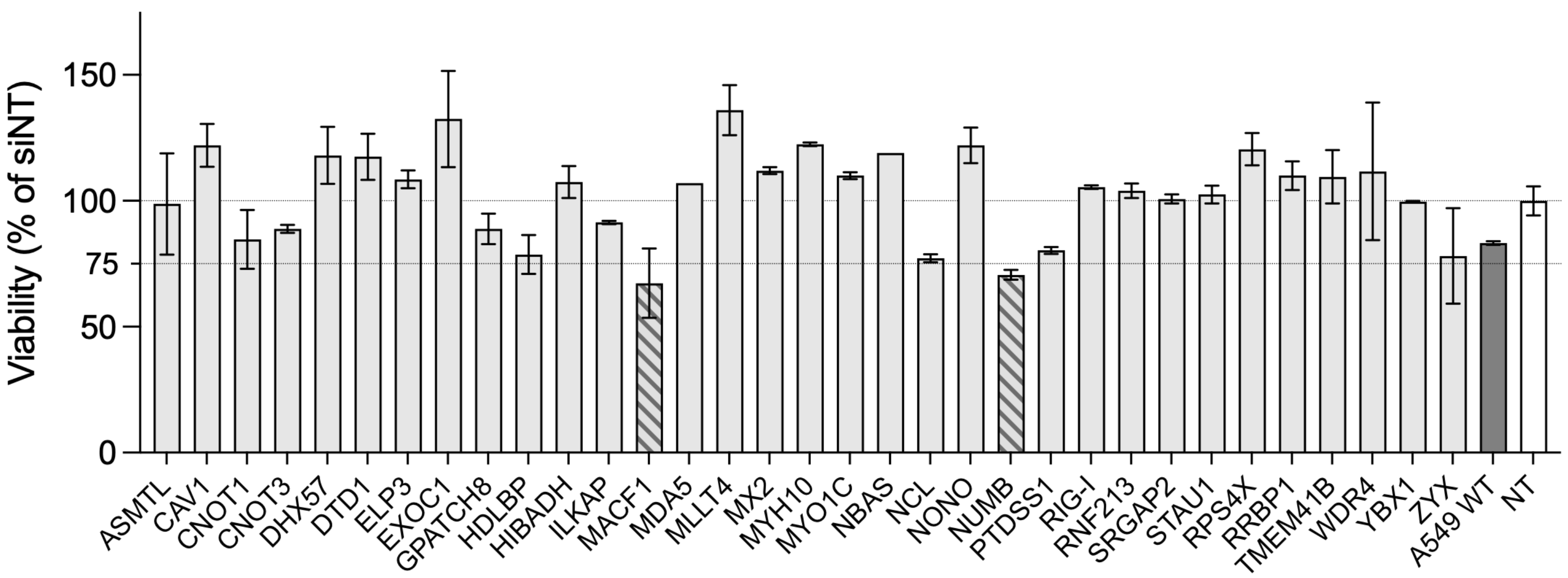


**B**

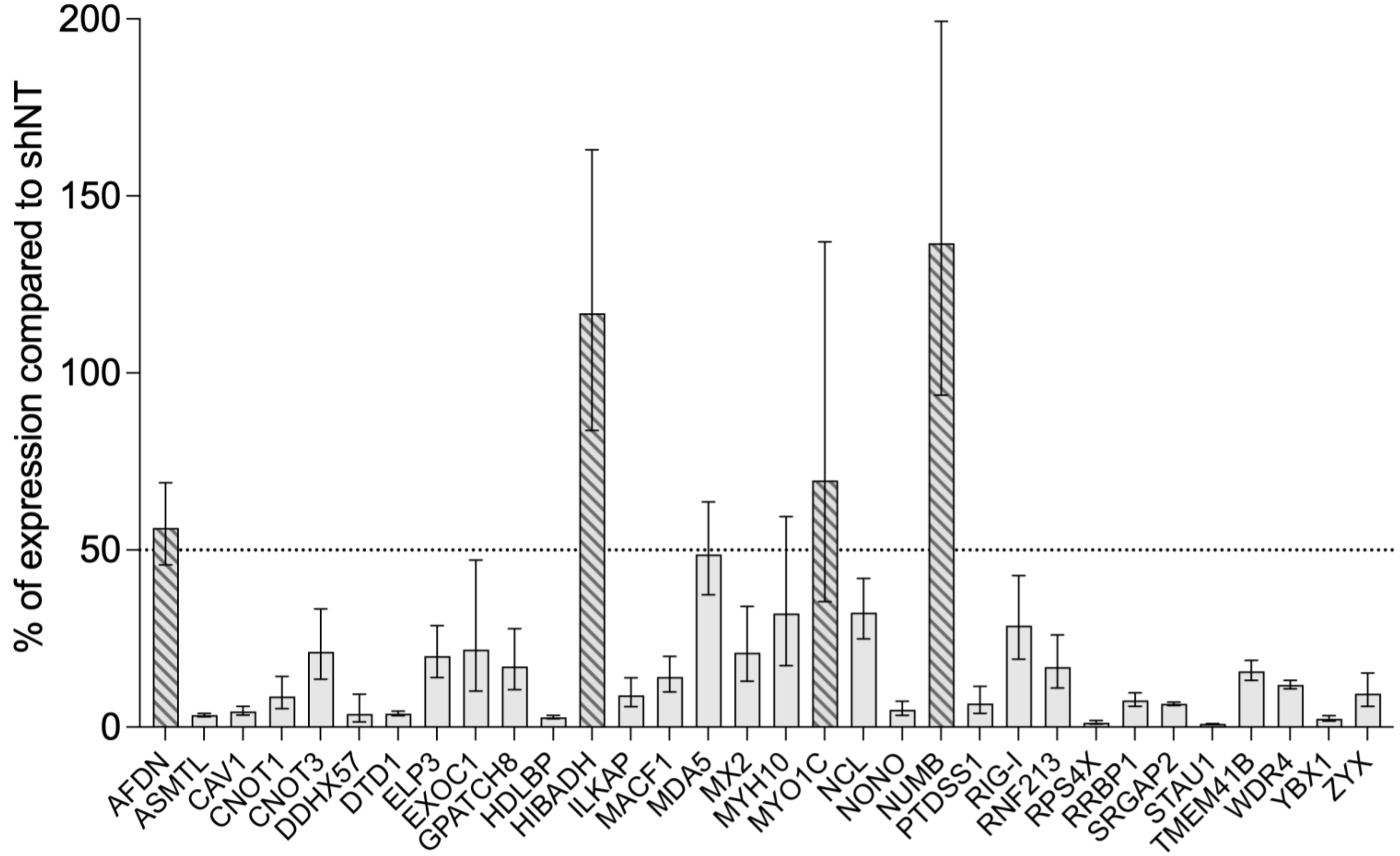


**Figure S2 - Viability and silencing efficiency of siRNA-treated cells**

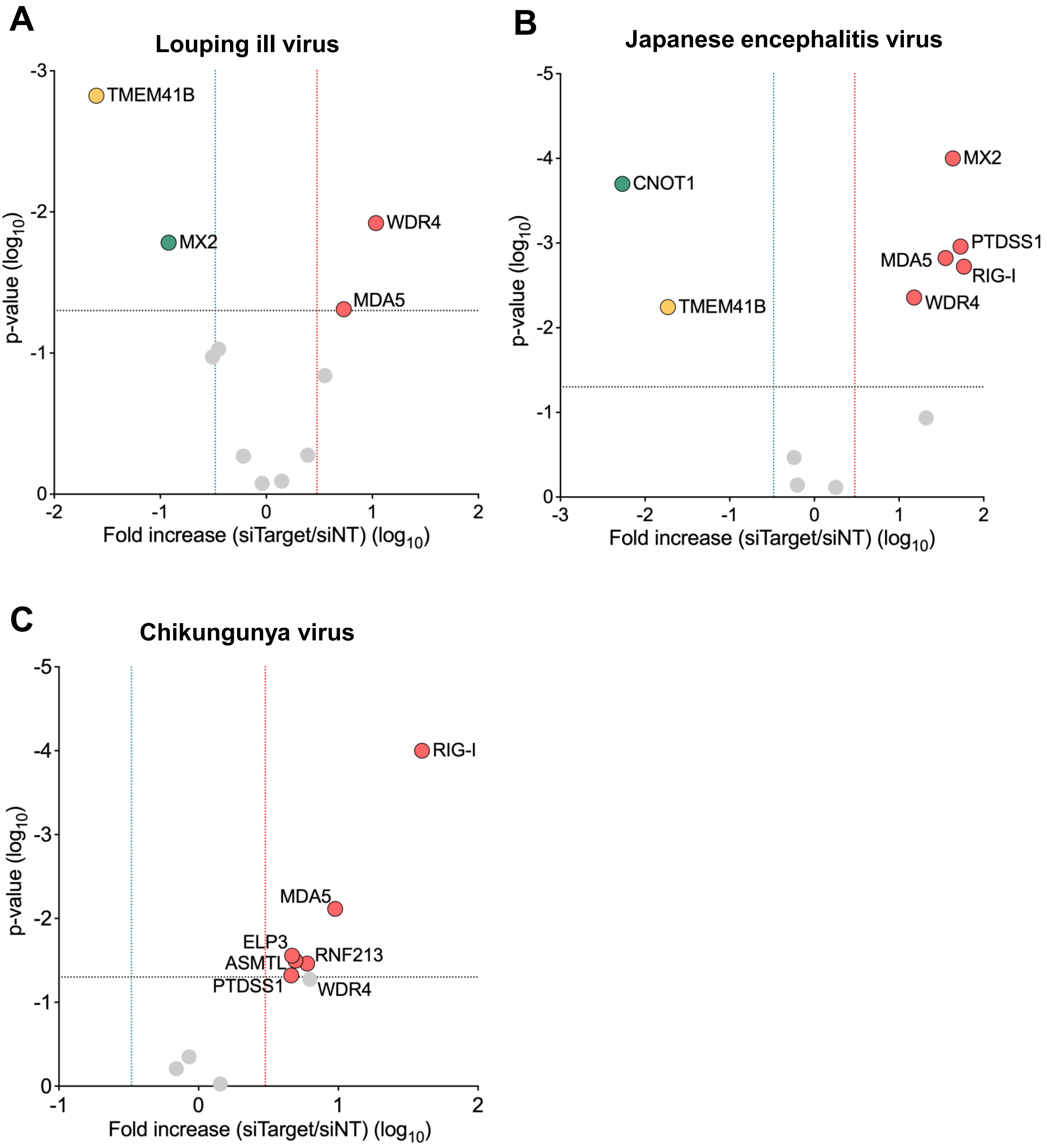


**Figure S3 - Functional analysis of impactful host factors for TBEV across diverse arboviruses**

**Figure S1 – Biological processes, molecular functions, and cellular compartments engaged by the viral RNA genome during TBEV infection**

Dendrograms depicting Gene Ontology enrichment for (A) biological processes, (B) molecular functions and (C) cellular compartments associated with the proteins in the extended interactome of the TBEV genome. GO-terms are clustered based on semantic similarity. Circles size and color reflect the number of enriched proteins within each term and the associated adjusted p-value, respectively. Only terms associated to an adjusted $p<0.05$ were retrieved and displayed.

**Figure S2 – Viability and silencing efficiency of siRNA-treated cells.**

(A) Cell viability and (B) gene-silencing efficiency were assessed in A549 cells transfected with siRNAs targeting the indicated host factors. All values are expressed relative to the siNT treated cells. Data represent the mean ± SEM of three biological replicates.

**Figure S3 – Functional analysis of impactful host factors for TBEV across diverse arboviruses**

Scatter plot showing the fold change in infectious viral particle release by siRNA silenced and (A) LIV-, (B) JEV- and (C) CHIKV-infected A549 cells, against statistical significance. Fold change is calculated as the mean viral titer in siTarget-transfected cells relative to siNT controls. Host factors are considered to significantly impact viral fitness if the titer ratio increases or decreases by at least 3-fold, with an adjusted p-value<0.05. Green, red, grey, and yellow dots represent proviral, antiviral, non-effective, and control proteins, respectively. Data represent means of 3 independent experiments, each performed in technical triplicates. Statistical significance was assessed using the non-parametric Kruskal–Wallis test, followed by Dunn’s multiple-comparison *post hoc* test with Benjamini–Krieger–Yekutieli two-stage step-up correction for multiple testing.
